# Regulatory Analysis of Root Architectural and Anatomical Adaptation to Nitrate and Ammonium in *Brachypodium distachyon*

**DOI:** 10.1101/2025.01.06.631519

**Authors:** Hamid Rouina, Dilkaran Singh, Christopher Arlt, Babak Malekian, Lukas Schreiber, Benjamin Stich, Amy Marshall-Colon

## Abstract

Root system architecture plays an important role in nitrate and ammonium uptake, the two primary nitrogen (N) forms essential for plant growth. Plants deploy different strategies to optimize the N uptake by roots, based on a complicated regulatory network that controls root phenotype and physiology. Here, we studied the response of root architecture to varying N applications in the model species *Brachypodium distachyon*. Using a combination of phenotypic and transcriptomic analyses, we examined how different forms and concentrations of ammonium and nitrate affect root growth, biomass allocation, and N uptake. N concentrations significantly influence root traits such as root length, root hair development, and aerenchyma formation in response to nitrate and ammonium. Plants grown in ammonium conditions had thin but highly branched roots, whereas nitrate application resulted in shorter, thicker roots with denser root hair at higher nitrate concentrations. Furthermore, using advanced co-expression network analysis, we identified an Atypical Aspartic Protease (APs) gene encoding an aspartyl protease family protein and a phosphoenolpyruvate carboxylase 1 (PEPC1) gene in brachypodium, which potentially control the root architectural and anatomical adaptions to different N form. APs expression showed a positive correlation with total root length and lateral root development, along with a negative correlation with root hair density. In contrast, PEPC1 exhibited positive correlations with cortex, stele, root cross-sectional areas, and root hair density, while showing a negative correlation with total root length. These genes likely play an important role in the transcriptional regulatory networks involved in these adaptive responses, which highlight the complex interplay between root morphology, nitrogen metabolism, and environmental nutrient conditions.

## Introduction

Nitrogen (N) uptake by plant roots is dependent on the interplay among root physiological, morphological, anatomical, and transcriptional phenotypes (Liu et al., 2015; Rogers & Benfey, 2015). The study of the genetic mechanisms underlying N uptake is challenging, partly due to the difficulty of root phenotyping caused by limited accessibility, as well as the influence of soil heterogeneity and other environmental factors (Comas et al., 2013; Pereira et al., 2021). However, the critical and adaptive roles of roots in responding to different N applications cannot be overlooked. Although N application responses vary among crop species, most of the root architectural and anatomical traits show genetically conserved responses (Jia & von Wirén, 2020). However, compared to the model plant Arabidopsis there is still a considerable gap in knowledge regarding these mechanisms in crop species, where most studies remain at the physiological level.

To maximize yield under unfavorable environmental conditions, plants employ dynamic resource allocation strategies as adaptive mechanisms in response to stress. Strategies to improve N uptake efficiency often suggest modifying root architecture, such as developing deeper roots or reducing crown root number in maize to optimize resource uptake under low N conditions (Mi et al., 2010; Saengwilai, Tian, & Lynch, 2014; Trachsel et al., 2013). In species like brachypodium and rice, research indicates that N availability influences root traits and biomass allocation (Jian-Bo et al., 2010; Poiré et al., 2014). In addition, in maize, research indicated that N deficiency enhances water and nutrient absorption efficiency, increases carbon allocation to roots and accelerates root growth (Gaudin et al., 2011; Xiaoli et al., 2020). Also, other anatomical adaptations, such as cortical aerenchyma and larger metaxylem, positively contribute to N acquisition by reducing metabolic costs and enhancing nutrient uptake (Klein et al., 2020; Saengwilai, Nord, et al., 2014). It has been demonstrated that the living cells within the cortex of the root segment are responsible for substantially affecting the maintenance cost of the root biomass in general (Jaramillo et al., 2013; Saengwilai, Nord, et al., 2014). The development of aerenchyma in roots, a response observed across several species like brachypodium, wheat, and maize, reduces metabolic demand and aids in nutrient acquisition under low N conditions (Drew et al., 1989; Khan et al., 2015; McDonald et al., 2002). This synergistic interaction between root architectural and anatomical characteristics is very important in the strategy for N acquisition (Galindo-Castañeda et al., 2022).

Studies of the root transcriptome in brachypodium as a model plant for monocot crops demonstrated that carbon metabolism in roots is tightly regulated to support N assimilation, with specific enzymes like phosphoenolpyruvate carboxylase (PEPC) playing crucial roles in balancing carbon and N metabolism (Caburatan & Park, 2021; de la Peña et al., 2019; Nimmo, 2006; Shi et al., 2015). PEPC activity, especially in roots, is integrated to metabolic processes, influencing growth and nutrient allocation (Chen et al., 2004; Masumoto et al., 2010; Shi et al., 2015). Shi et al. (2015) demonstrated that the other isoform of PEPC, PPC3, which is found highly expressed in roots, plays a significant role in root metabolism. It has also been shown to play a key role in malate production in root nodules and could thus be responsible for N metabolism via malate synthesis in the Arabidopsis root (Chen et al., 2004). Several studies have shown evidence that the addition of nitrate and ammonium has different effects on PEPC and therefore affect carbon metabolism and allocation in roots (Koga & Ikeda, 1997; Pasqualini et al., 2001; Prinsi & Espen, 2018). However, the association between PEPC and various root traits is unknown, especially in the context of soil N availability.

Hormonal interactions also govern root development under varying environmental conditions. Auxin, a key plant hormone (Mazzoni-Putman et al., 2021), integrates developmental cues with environmental signals and influences root architecture (Du & Scheres, 2018; Tognetti et al., 2012). The overexpression of Atypical Aspartic Protease in Roots 1 (ASPR1) was shown to disrupt auxin balance and affected root growth negatively (Soares et al., 2019). Several studies suggested that atypical APs have specialized roles in plant physiology, indicating potential targets for enhancing root development under nutrient stress (Prasad et al., 2010; Simoes & Faro, 2004; Van Der Hoorn, 2008). It has also been demonstrated that ASPR1 plays a significant role in root development in Arabidopsis and maize. Its expression leads to shorter primary roots and reduced lateral root formation, indicating that ASPR1 negatively regulates these growth processes in plants (Liu et al., 2023; Soares et al., 2019). Atypical Aspartic Proteases may have broader functions in root development, potentially acting as key regulators involved in hormone signaling and root growth regulation. Nevertheless, only a limited number of studies have demonstrated the atypical role of APs in plant root development, and further research is necessary to examine its function in plant root traits. Specifically, the regulatory roles of ASPR1 and other atypical APs in interacting with hormonal pathways to control root development under varying environmental conditions remain unclear.

To address these knowledge gaps, we used digital phenomics approaches that enable non-destructive phenotyping of root growth in combination with transcriptional analysis. Digital phenomics provide greater access to the root to identify its function and adaptation in varying nutrient conditions (Furbank & Tester, 2011). The combination of rapid, non-invasive phenotyping and transcriptome analyses offers a promising opportunity to advance our understanding of nutrient uptake physiology and genetics. When this approach is combined with the genomic tools available for *Brachypodium distachyon*, it has the potential to accelerate the identification of genes controlling N uptake and root architecture. We established the dose-dependent response of various root architectural and anatomical traits to ammonium and nitrate concentrations in the growth medium. Using RNA-seq on root samples from the lowest (0 mM), middle (1.5 mM), and highest (6 mM) N concentrations, we first identified genes correlated with general phenotypic responses to N availability. Subsequently, through Weighted Gene Co-expression Network Analysis (WGCNA), we explored groups of co-expressed genes, which specifically correlated with reduced root metabolic costs, as well as to distinct anatomical and morphological traits. This comprehensive approach uniquely facilitated the exploration of genes linked to both general phenotypic responses and specific traits, providing insights into the genetic and physiological mechanisms underlying root adaption in response to N application.

## Results

### Phenotypic analysis

#### Ammonium and nitrate treatments affect biomass allocation and N uptake

We evaluated the effect of N form and concentration on biomass allocation between roots and shoots, as well as overall N uptake. Whole plants were harvested to measure N content, as well as fresh and dry weights. The total dry mass increased as N concentration increased; however, ammonium treated plants showing significantly greater biomass (P value =2.14×10^-3^) than those receiving nitrate (Fig-1A). This increase in total dry mass was primarily due to increased shoot biomass, while root biomass decreased with increasing N concentration (Fig-1A). At lower N concentrations, around 65% of total biomass was allocated to the roots, while this allocation dropped to approximately 30% at higher N concentrations. Across all N concentrations, plants grown with ammonium generally took up more N per milligram dry weight than those grown with nitrate (Fig-1B), which likely contributed to the greater overall biomass of ammonium treated plants; a difference that became more evident in the higher range of N concentrations. The N content in plant tissues increased up to a concentration of 1.5 mM for both N forms and remained constant from 1.5 to 6 mM. However, nitrate treated plants absorbed more N per unit of root length, primarily because they achieved this uptake with a shorter total root length (Suppl 1).

**Figure 1:**
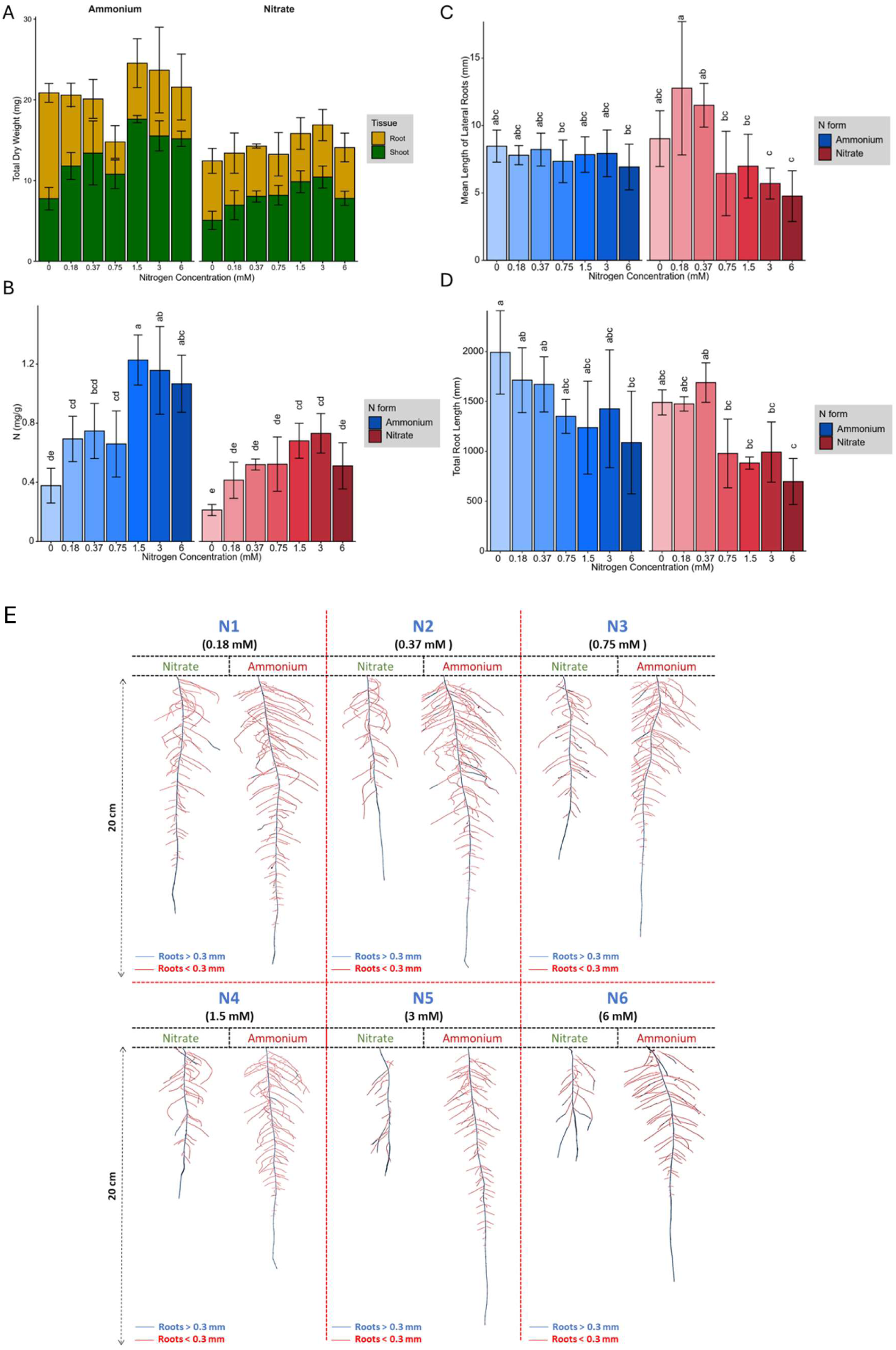
**(A)** Root and shoot dry biomass, **(B)** Nitrogen content per gram of total dry biomass, **(C)** Mean length of lateral roots (mm), **(D)** Total root length (mm), in response to 0 mM, 0.18 mM, 0.37 mM, 0.75 mM, 1.5 mM, 3 mM, and 6 mM ammonium and nitrate concentrations in the growth media. **(E)** Response of *Brachypodium distachyon* root systems architecture of plants grown under various nitrate and ammonium concentrations. The red color represents roots with a diameter of less than 0.3 mm (lateral roots), while the blue color represents roots with a diameter of more than 0.3 mm (seminal root). Image of representative plants in each condition. In (a) and (B) error bars represent standard error of mean, and in (B) letters indicate statistical significance between groups (p < 0.05).

#### Root system architecture (RSA) has differential responses to N forms and concentrations

Similar to root dry mass, total root length decreased with increasing N concentrations, where root architecture adapted to both N forms and concentrations (Fig-1E). Reduced total root length in response to increasing N concentration was accompanied by decreases in root system area and root volume (Suppl 2). However, compared to nitrate-grown plants, ammonium-grown plants exhibited greater total root length, network area, root volume, and surface area with less variation in root phenotype across different total N concentrations. In contrast, plants grown with nitrate exhibited two distinct response plateaus, one at low N concentrations, corresponding to the range of high-affinity nitrate transporters (HATS) (0 to 0.37 mM), and another at higher concentrations, associated with low-affinity nitrate transporters (LATS) (0.75 mM to 6 mM) (Fig-1D). Across all treatments, each plant produced only one seminal root, which was longest under low N conditions, with its length decreasing as N concentrations increased. This decline was particularly pronounced with nitrate application. Longer seminal roots produced more lateral branches (Suppl 2), although branching frequency remained relatively constant across different N concentrations and sources (Suppl 3).

#### N forms and concentrations shape root hair development and lateral root architecture

Plants treated with ammonium developed a more extensive root area primarily due to an increase in the number of branched roots. Although low nitrate concentrations also stimulated more root branching, the average length of lateral roots remained relatively stable across the different N concentrations and forms (Fig-1C). However, at lower nitrate concentrations (0.1875 mM and 0.375 mM), plants developed significantly longer lateral roots. In contrast, at higher N concentrations (1.5 mM, 3 mM, and 6 mM), ammonium treated plants exhibited a significantly greater total lateral root length compared to those grown in nitrate (Pvalue= 7.28×10^−9^). Additionally, ammonium treated plants produced more secondary-order lateral roots, whereas increasing nitrate concentrations induced the formation of root hairs (Fig-2A). Microscopic examination revealed that the formation of root hairs on seminal roots was significantly (Pvalue= 2×10^−16^) affected by both the form and concentration of N. Plants exposed to ammonium had reduced root hair density and length (Fig-2B). In contrast, plants treated with nitrate showed distinct responses depending on the concentration. At low nitrate concentrations, root hairs were longer, but had significantly lower density compared to those at higher nitrate concentrations (Pvalue= 2×10^−16^), where root hair density increased despite a reduction in their length (Fig-2C).

**Figure 2:**
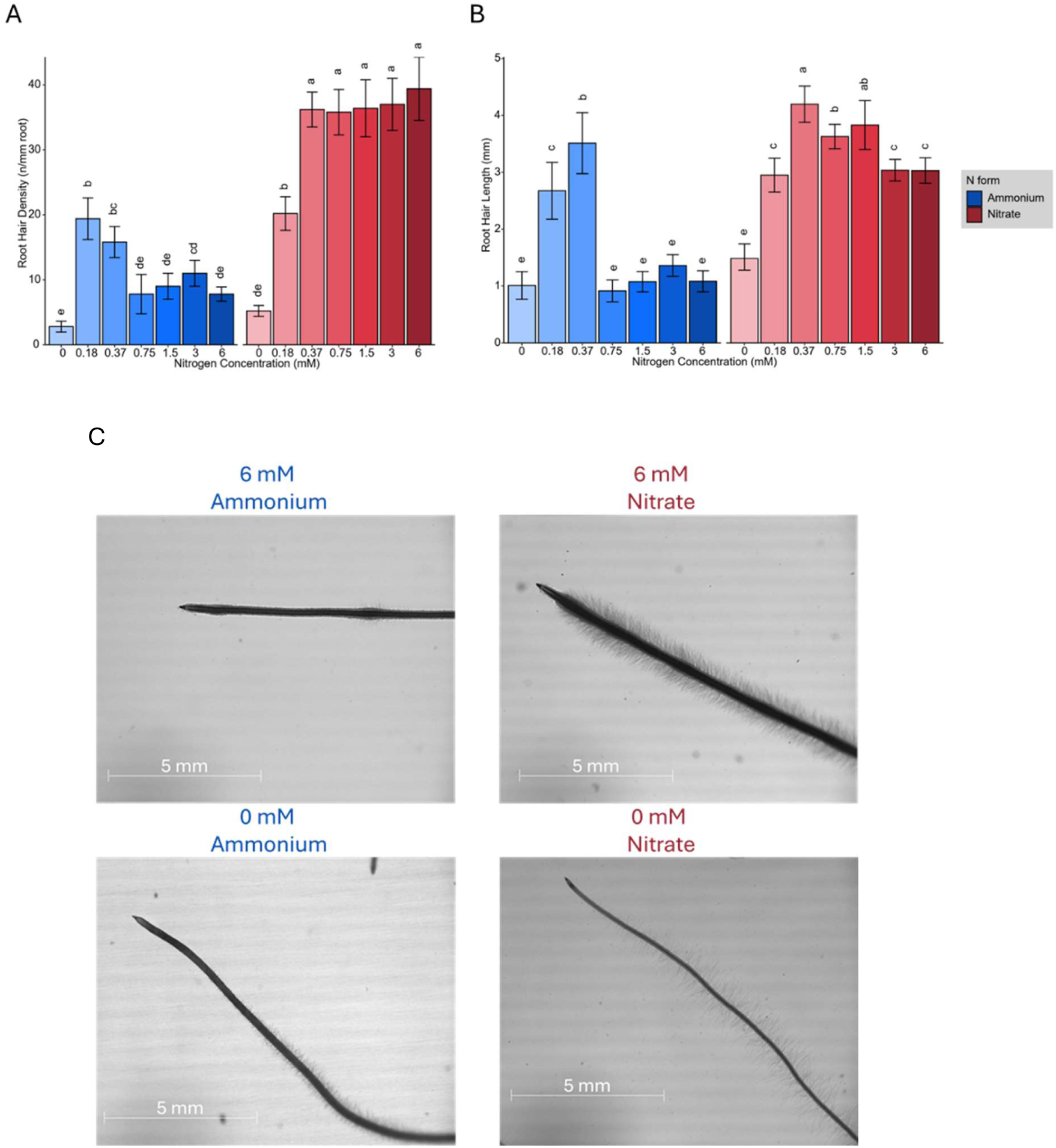
**(A)** Root hair density (hairs per mm root), **(B)** Root hair length (mm) of *Brachypodium distachyon* plants grown under diferent nitrogen concentrations (0, 0.18, 0.37, 0.75, 1.5, 3, and 6 mM) of ammonium (blue) and nitrate (red). **(C)** Representative images of brachypodium root hair morphology at 6 mM and 0 mM of ammonium (left) and nitrate (right). Scale bars represent 5 mm. in (A), (B), (C), and (D) error bars represent standard error of mean, and letters indicate statistical significance between groups (p < 0.05).

#### Root aerenchyma formation and anatomical features have distinct differences in response to ammonium and nitrate

We examined the effects of varying N concentrations on root anatomy by dividing the seminal roots into three sections of equal-length and taking cross-sections of each from root base, middle, and tip (suppl 4). The anatomy of the roots varied significantly (Pvalue= 5.47×10^−4^) across these zones (Fig-3E). The root cross-sectional area increased with rising nitrate concentrations. At high nitrate concentrations, the root cross-sectional area was larger in the younger root sections compared to the basal sections (Fig-3A). In contrast, ammonium application significantly (Pvalue= 2.66×10^−11^) reduced the cross-sectional area, with younger root sections being smaller (Fig-3A). Thicker roots with greater cross-sectional areas generally exhibited an expanded cortical area without a corresponding increase in stele area (Fig-3B). This expansion of the cortical area was primarily due to an increase in cell size rather than cell number (suppl 11). Although the stele area remained relatively unchanged (suppl 11), the central metaxylem area measured as fraction of the total stele area, significantly increased (Pvalue= 2.27×10^−7^) from the root base to the middle and tip in both ammonium and nitrate treatments across all concentrations (Fig-3C). Ammonium treated roots were generally thinner than nitrate treated roots, primarily due to a reduced cortex area (Fig-3E). Higher ammonium concentrations not only decreased cortical thickness but also induced the formation of aerenchyma in the upper (older) sections of the roots (Fig-3D). Consequently, the number of cortical cells decreased. Plants exposed to high ammonium concentrations had a smaller living cortical area, resulting from a combination of reduced cortical cell width and fewer living cortical cells. Notably, the basal sections of the roots did not show significant differences in most measured traits in response to either N concentration or form.

**Figure 3:**
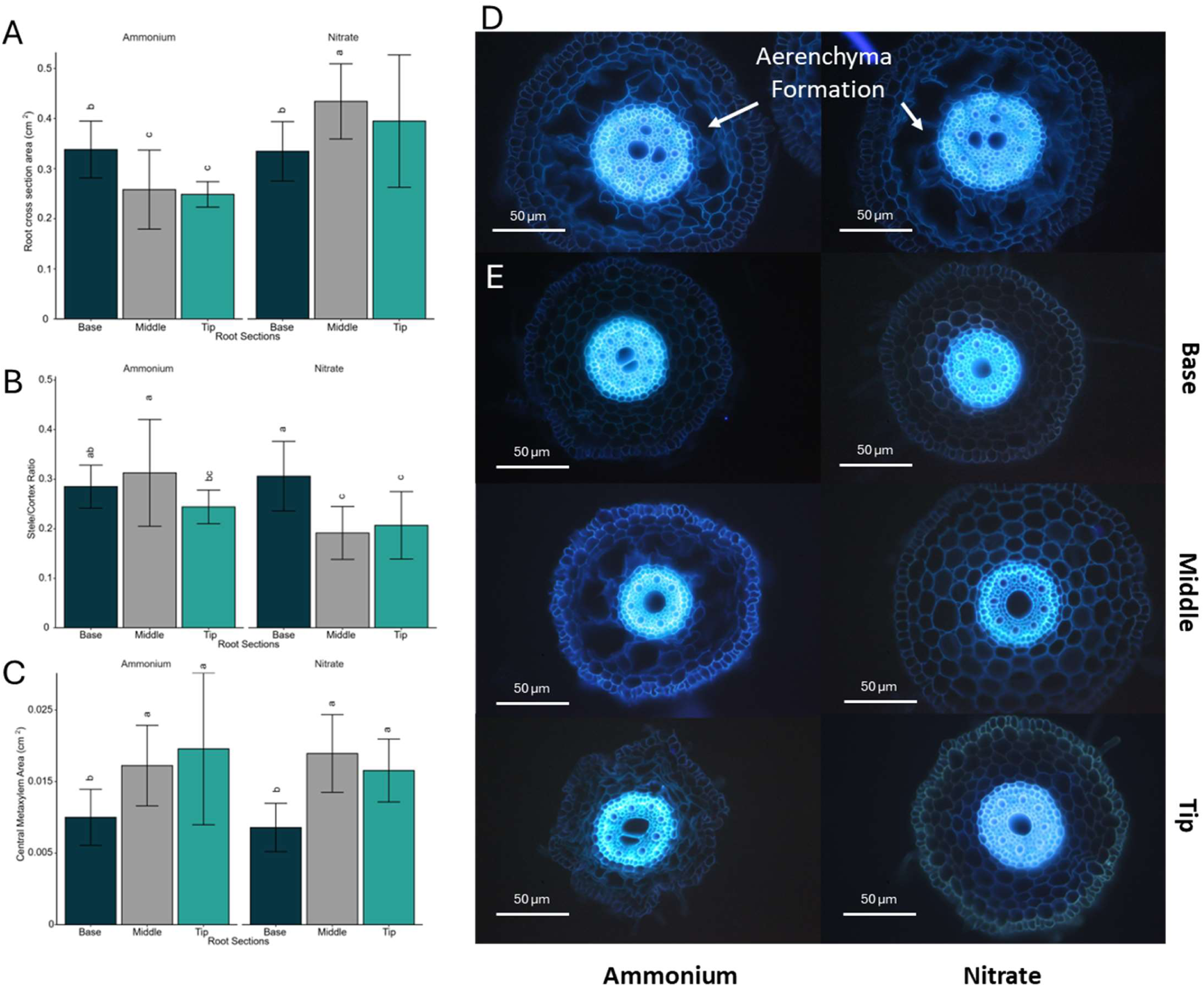
Root anatomical characteristics assessed in their response to ammonium and nitrate. **(A)** Root cross-sectional area (cm²), **(B)** Stele/cortex ratio, **(C)** Central meta xylem area (cm²) in three regions of the root (Base, Middle, Tip) (dark green /gray /green) under ammonium (left) and nitrate (right) treatments. **(D, E)** Fluorescence microscopy images of root cross-sections show structural diferences between ammonium (left) and nitrate (right) treatments. The sections are arranged by root region (Base, Middle, Tip). in (A), (B), and (C) error bars represent standard error of mean, and letters indicate statistical significance between groups (p < 0.05).

### Transcriptional analysis

#### Nitrogen-responsive genes are differentially expressed in response to N form and concentration

Using a generalized linear model (GLM) analysis across all comparisons of N concentrations we identified 5,517 genes that were significantly differentially expressed under ammonium application, and 2,538 genes that were significantly expressed in response to nitrate applications (suppl 5). For both N forms, we used the lowest N concentration as the reference and compared all other N concentrations against it. Most of these genes were differentially abundant between high and low N concentrations, with few significant differences observed between low and moderate or moderate and high concentrations (suppl 5). The clustering patterns observed in the Principal Component Analysis (PCA) of transcript levels closely mirrored those from Root System Architecture (RSA) measurements, which were based on 29 distinct traits (Fig-4). The PCA consistently grouped samples from the same treatment together, indicating a strong alignment between gene expression profiles and root system characteristics (Fig-4).

**Figure 4:**
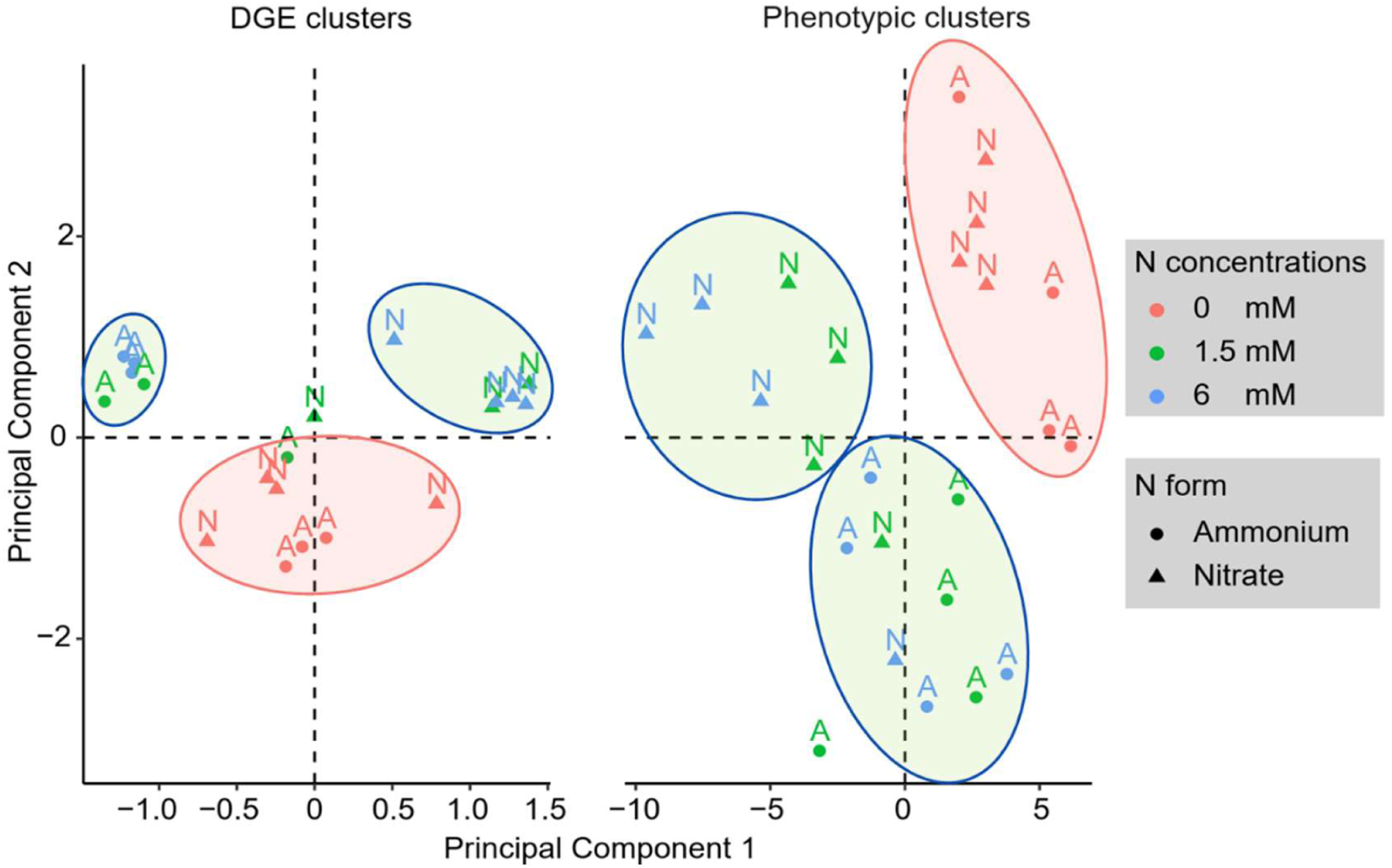
Principal component analysis of *Brachypodium distachyon* gene expression (left) and phenotypic traits (right). The scatter plots illustrate the clustering of diferentially expressed genes (DGE clusters) and phenotypic traits (Phenotypic clusters) under varying nitrogen levels (0, 1.5, and 6 mM) and nitrogen sources (ammonium and nitrate). Color coding reflects nitrogen concentrations (red for 0 mM, green for 1.5 mM, and blue for 6 mM), while the shape represents nitrogen form. Clusters are visually separated by ellipses to indicate groupings of similar responses.

To visualize the biological processes affected by N treatment, the gene ontology of significantly expressed genes were found using the “enricher” function of the “Biomart” library in R (Durinck et al., 2009). Gene ontology (GO 2.62.0) enrichment analysis revealed distinct patterns in gene regulation in response to N application. In nitrate treated plants, genes involved in the response to other nutrients (such as phosphate, cadmium, and sulfate) were upregulated in both high and moderate nitrate conditions compared to low nitrate conditions. Conversely, gene families associated with N transport were downregulated at moderate nitrate concentrations compared to low nitrate concentrations (suppl 6). In ammonium treated plants, high ammonium concentrations led to upregulation of genes related to cytokinin activity and chloroplast functions. However, genes associated with auxin activity, carbohydrate transport, and water channels were downregulated at both high and moderate ammonium concentrations compared to low ammonium concentrations. Notably, genes involved in anatomical development and cellulose activity were also downregulated at high ammonium concentrations suggesting their close relationship with ammonium uptake and metabolism (suppl 7).

#### High and low affinity nitrate transporter and ammonium transporter activities respond to N applications

Overall, the gene expression/transcript levels of nitrate and ammonium transporters was more sensitive to N concentration than to N form. In brachypodium, five genes identified as low-affinity nitrate transporters were analyzed separately (suppl 8) (Guo et al., 2014; Sigalas et al., 2024). The differentially expressed genes (DEGs) analysis revealed that the NRT1.1 gene, a low-affinity nitrate transporter, was upregulated in response to higher nitrate applications (Fig-5). However, NRT1.2 and NRT1.3 did not show significant (Supplementary data 1) changes across nitrate concentrations (Fig-5). Conversely, NRT1.4 and NRT1.5 were upregulated at moderate nitrate concentrations (Fig-5) compared to low nitrate concentrations. Among high-affinity nitrate transporters, seven out of nine members were expressed in both ammonium and nitrate application, with five belonging to the NRT2 family and two to the NRT3 family. The NRT3 gene family showed consistent expression across all nitrate concentrations (Fig-5). In contrast, NRT2.1, NRT2.2, and NRT2.4 were upregulated with increasing nitrate concentrations, while NRT2.5 and NRT2.7 were dramatically (Supplementary data 1) downregulated at high nitrate concentrations (Fig-5). Additionally, at high ammonium concentrations, the gene families responsible for ammonium transporters (BRADI_1g02420v3 and BRADI_3g45480v3) were downregulated by approximately two-fold (Fig-5).

**Figure 5:**
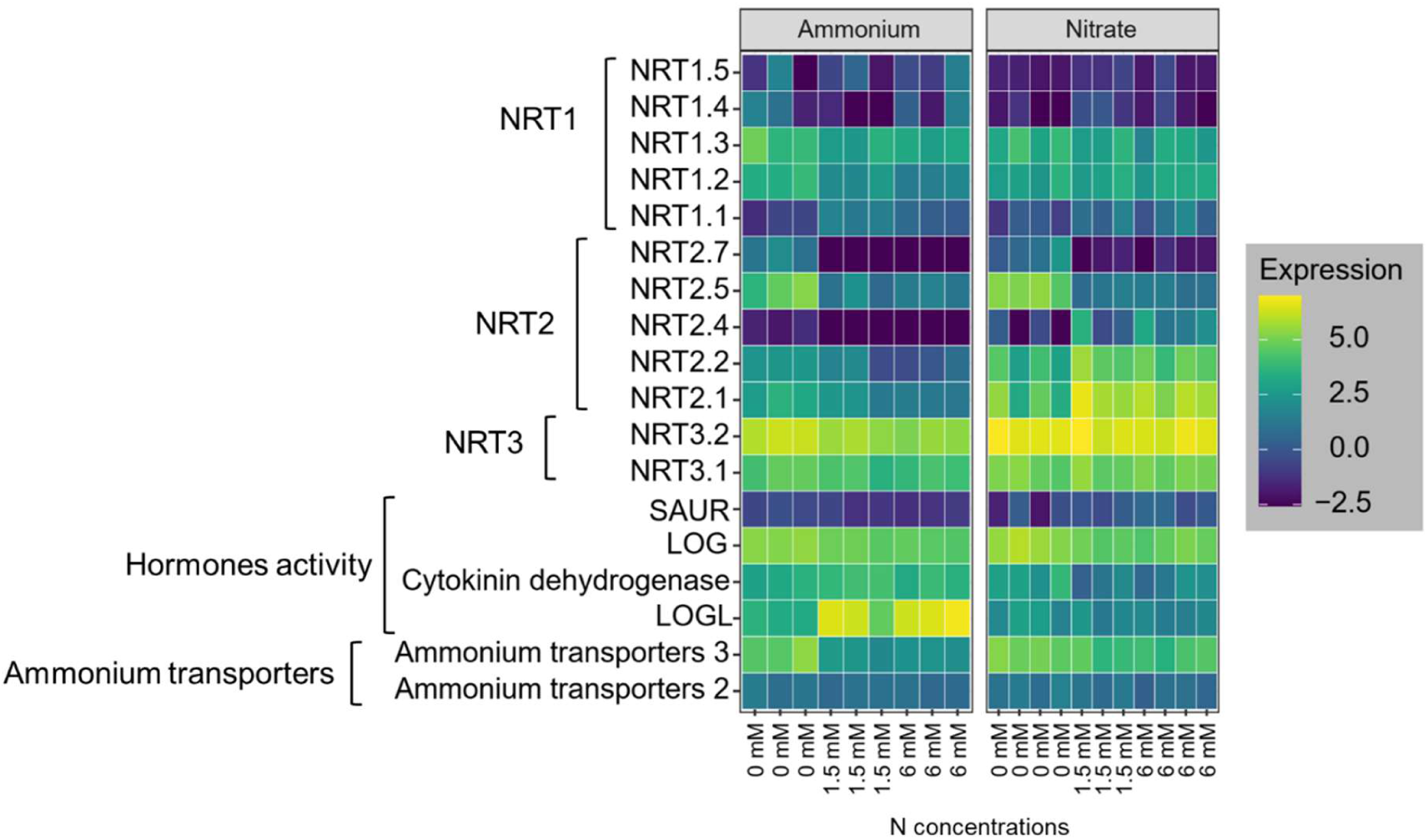
Heatmap of gene expression levels related to nitrate transporters (NRT1, NRT2, and NRT3), hormones activity, and ammonium transporters in brachypodium roots under ammonium and nitrate treatments at varying nitrogen concentrations (0, 1.5, and 6 mM). The experiment had three replicates.

#### Contrasting phytohormone activity was observed in response to ammonium and nitrate

Gene ontology enrichment analysis revealed significant changes in phytohormone activity in plants grown with ammonium compare with those in nitrate. Specifically, genes associated with cytokinin, and auxin pathways were notably affected. In ammonium-treated plants, a downregulation of genes involved in auxin signaling, including the SAUR family gene BRADI3g05020v3 (Fig-5) was observed. Conversely, the cytokinin-related gene LOGL (BRADI3g28900v3) was significantly upregulated in ammonium-treated roots (Pvalue= 1.66×10^-6^ for Low ammonium vs moderate ammonium, Pvalue= 1.75×10^-7^ for Low ammonium vs high ammonium), suggesting its role in promoting lateral root growth and development (Fig-5). However, some other cytokinin signaling genes, including LOG (BRADI2g42190v3) and Cytokinin dehydrogenase (BRADI2g05580v3), were downregulated in response to both ammonium and nitrate applications regardless of their concentration (Fig-5). This targeted analysis of hormonal response genes helps clarify the molecular mechanisms by which different nitrogen forms modulate root development, as indicated in prior studies (Asim et al., 2020; Dziewit et al., 2024; Kazan, 2013; Naulin et al., 2019).

#### Aquaporin related genes and water channel activity in ammonium application

Given the crucial role of aquaporins in regulating water and nitrogen transport within plant roots, analyzing specific aquaporin gene responses under varying N sources can help elucidate their roles in managing nutrient uptake and preventing nitrogen toxicity (Wang et al., 2016). The aquaporin gene family members TIP4-2 (BRADI_2g07830v3), TIP2-1 (BRADI_2g62520v3), TIP5-1 (BRADI_5g17680v3), and TIP4-3 (BRADI_2g07810v3) were downregulated in plant roots grown under elevated ammonium concentrations (Suppl 9). Conversely, in roots grown under nitrate conditions, these genes (except TIP4-2) were upregulated as the N concentration increased. The downregulation of TIP4-2 in response to higher ammonium concentrations aligns with the downregulation of ammonium transporters, suggesting a similar role in preventing N excessive accumulation, particularly as total N uptake per unit root length becomes saturated at 1.5 mM external N (Fig-1B).

### Integrated phenotypic and transcriptomic analysis

#### Correlation of gene modules with root traits highlights distinct N uptake strategies

To uncover key genes contributing to the observed root phenotypes upon N treatment, a Weighted Gene Co-expression Network Analysis (WGCNA) was performed across ammonium and nitrate treatments. This approach identified 32 co-expressed gene modules across various ammonium and nitrate concentrations, and clusters were distinguished by different colors in the hierarchical clustering dendrogram (Fig-6). Each module contained from 45 to 2,923 genes. We tested the correlation of each module’s eigengene with the root morphological and anatomical phenotypes as well as N content of the plants. Correlation analysis of module eigengenes with phenotypic traits revealed several significant associations (Fig-7).

**Figure 6:**
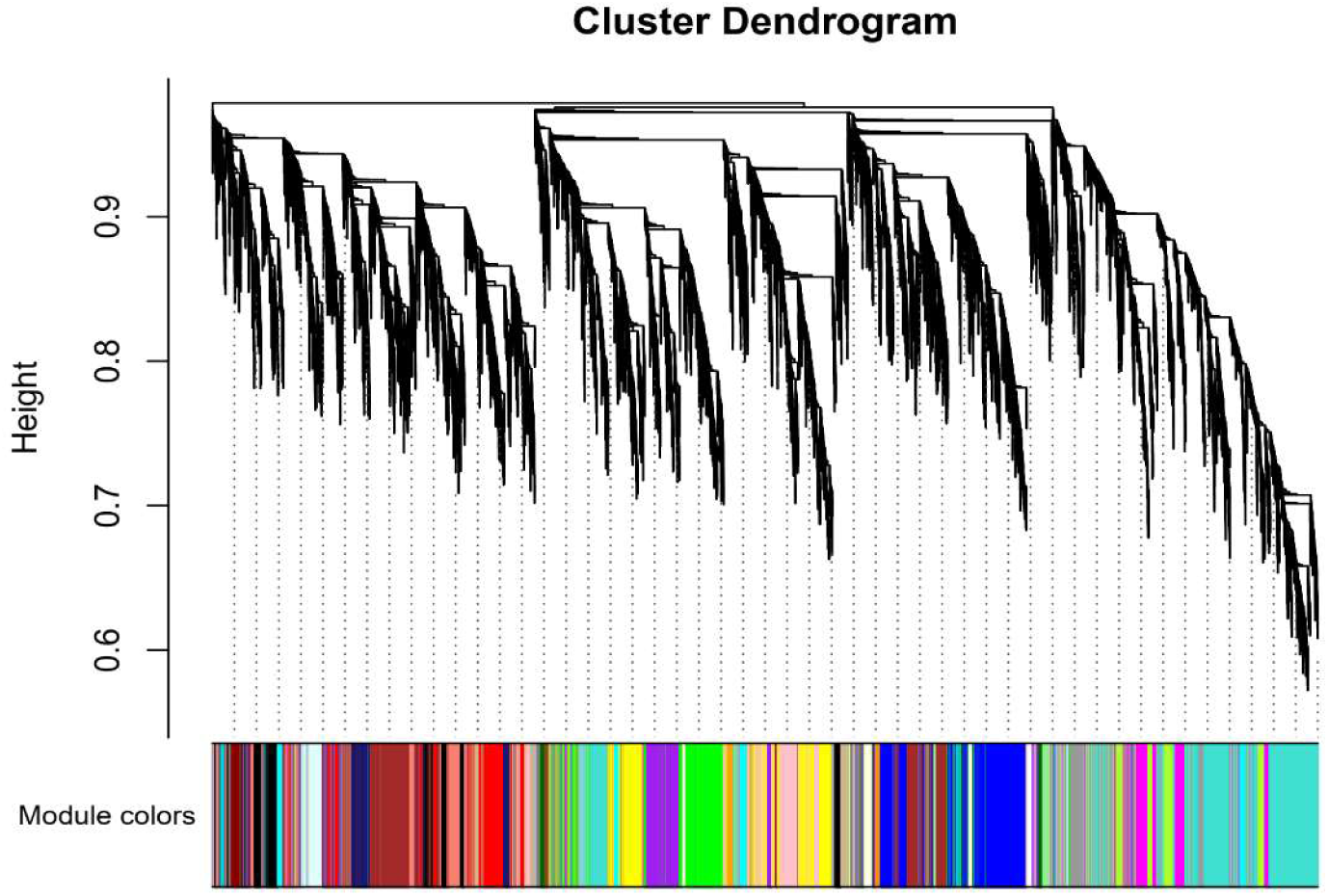
The dendrogram shows the hierarchical clustering of genes into modules based on their co-expression patterns. Each module is represented by a diferent colored bar below the dendrogram. Modules group genes that are highly co-expressed, indicating similar functional or regulatory roles in response to nitrogen treatments.

**Figure 7:**
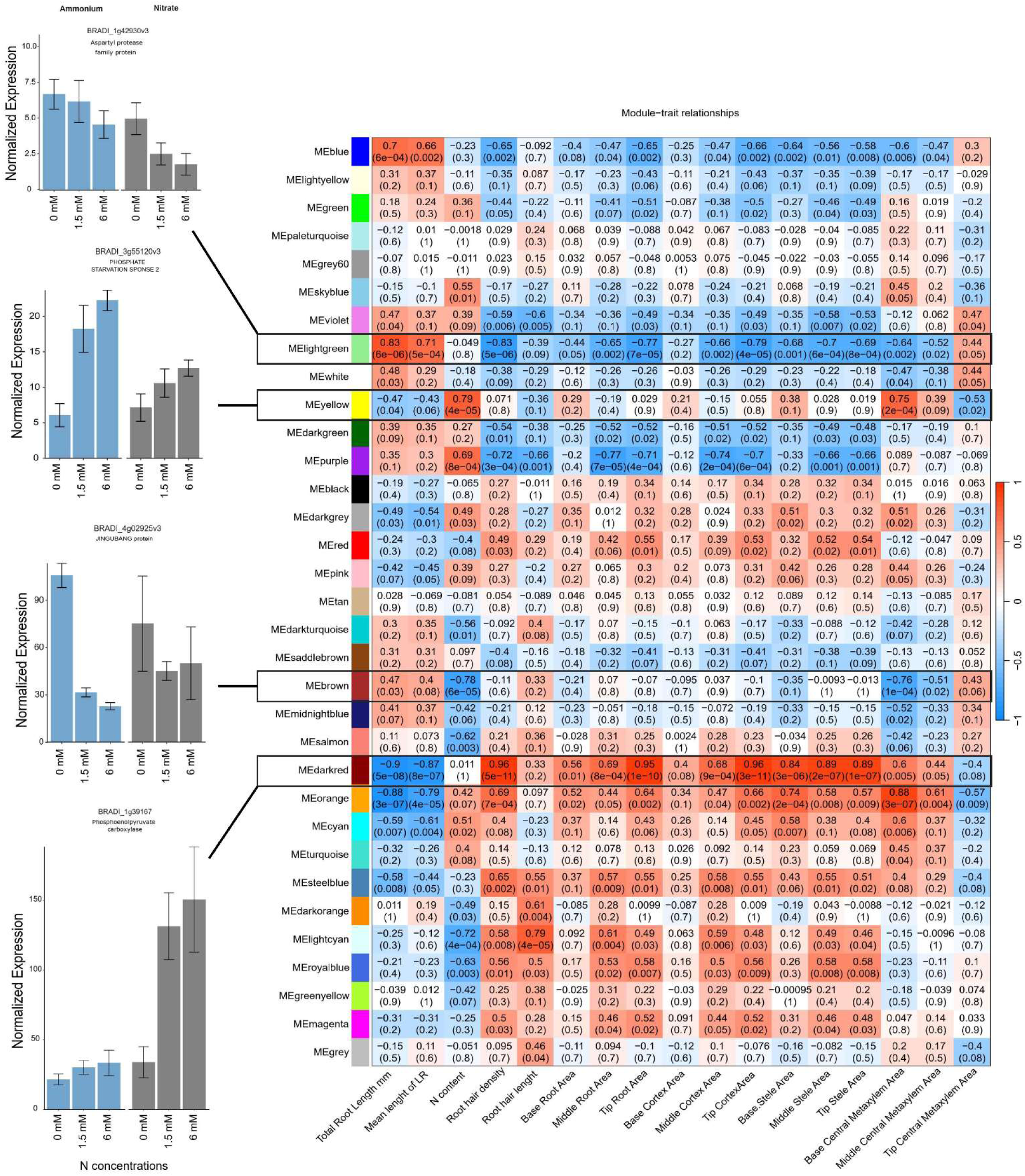
The heatmap (right) depicts the correlation between gene expression modules and root traits of interest across nitrogen treatments. Each cell represents the Pearson correlation coeficient between a module’s eigengene (the first principal component of gene expression within a module) and a specific trait, with corresponding p-values in parentheses. Positive correlations are shown in blue and negative correlations are shown in red. Bar plots (left) are representative of each module normalized gene expression in *Brachypodium distachyon* in response to nitrogen-free, 1.5 mM, and 6 mM ammonium (left) and nitrate (right) conditions. Error bars represent the standard error of the mean, with normalization performed using the TMM method.

Two modules, “light green” (including 350 genes) and “dark red” (including 205 genes), exhibited striking contrasts with root traits (Supplementary data 4). The “light green” module was positively correlated with traits related to root elongation (total root length (|r| = 0.83) and mean lateral root length (|r| = 0.71)), and negatively correlated with root hair traits and anatomical features (Fig-7). In contrast, the “dark red” module was associated with more compact root systems, but it showed positive correlations with root hair development (root hair length |r| = 0.33 and density |r| = 0.96) (Fig-7). These contrasting gene-to-trait associations between the two modules demonstrates different strategies of root adaptation to N availability, either elongation to explore more soil volume (“light green”) or increasing surface area through root hairs (“dark red”). The “yellow” (including 1619 genes) and “brown” (including 1770 genes) modules also displayed contrasting relationships with respect to N content (Supplementary data 4). The “yellow” module was positively correlated with total N content (|r| = 0.79), while the “brown” module was strongly negatively correlated with N content (|r| = 0.78), suggesting different roles for the genes within these modules toward N uptake or storage efficiency (Fig-7).

#### Intramodular analysis identifies key genes correlated with root phenotypes

For each module, genes with high Gene Significance (0.9 < GS) and Module Membership (0.8<MM) scores were examined as potential key genes for the corresponding phenotypes. This intramodular analysis enabled us to identify genes that may contribute to the underlying mechanisms governing root development in response to N application. We identified key genes within the modules that showed significant correlations with various root phenotypes. Through ontology enrichment of these genes, and their homologues in rice, Arabidopsis, and maize, we elucidated their function using the gProfiler (Raudvere et al., 2019) and NCBI database for annotation, visualization, and integrated discovery (DAVID) tool (Huang et al., 2007). Among these genes, we selected genes with top rank and significant expression for further analysis (Table 1).

**Table1:**
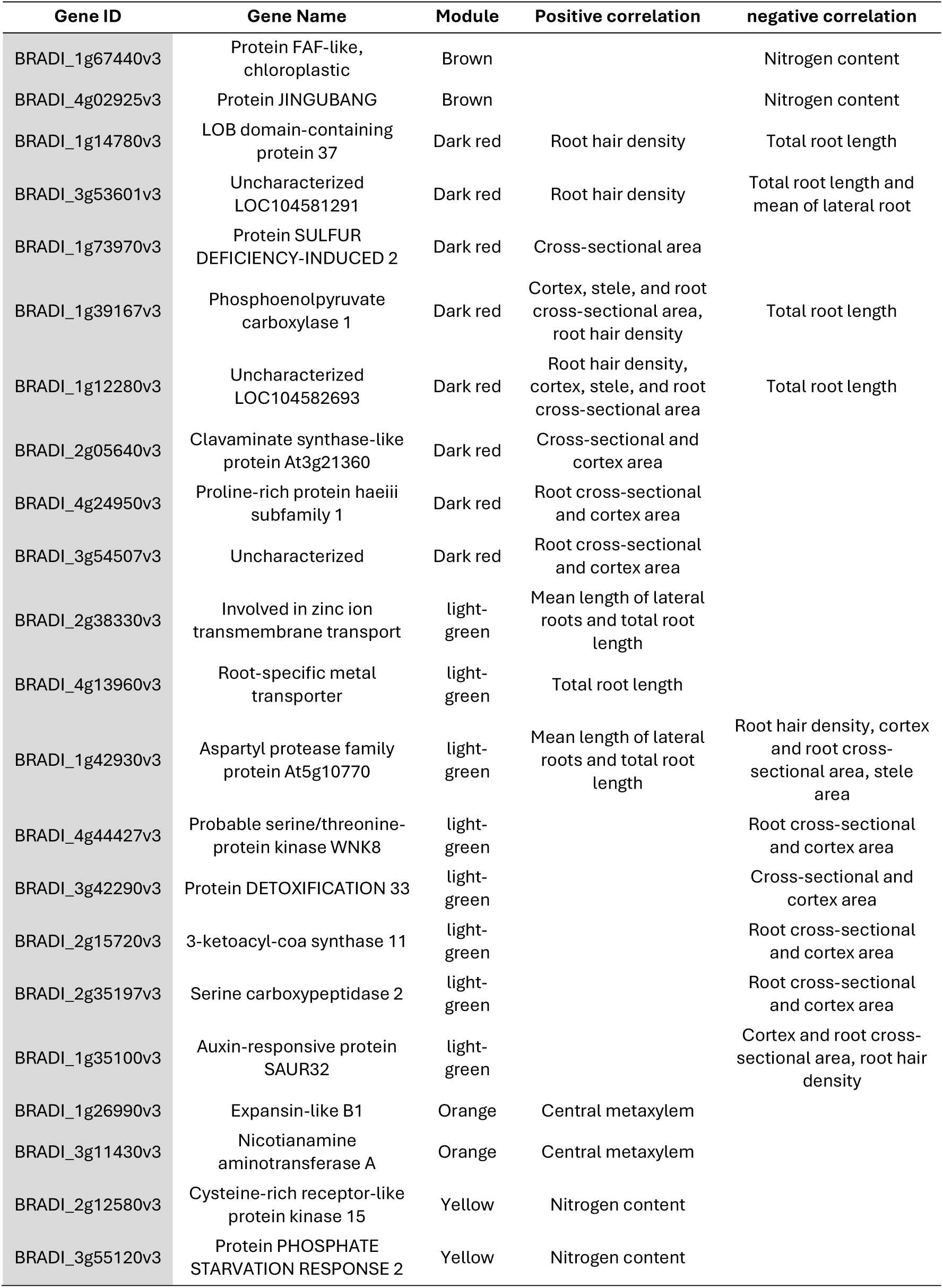
Genes identified with high gene significance and module membership scores in each module for root phenotypes in response to N application.

Following these criteria, investigation of the light green module identified the potential hub genes BRADI_1g42930v3, a homolog of an aspartyl protease family protein (APs, At5g10770), BRADI_1g35100v3 (SAUR32), a gene encoding an auxin-responsive protein, BRADI_2g38330v3, which is involved in zinc ion transmembrane transport, and BRADI_4g13960v3, a gene associated with root-specific metal transporter responses. These genes had similar expression patterns under low N conditions but were downregulated under high N conditions, with more pronounced suppression in the presence of nitrate than ammonium (Fig-7). As mentioned above, the light green module had strong positive correlations with the mean length of lateral roots and total root length but a strong negative correlation with root hair density in response to N concentration.

Investigation of the dark red module identified the hub genes BRADI_1g12280v3, homolog of HPP in maize and Arabidopsis, BRADI_1g39167v3 (phosphoenolpyruvate carboxylase 1, PEPC1), BRADI_1g14780v3 (LOB domain-containing protein 37), and BRADI_1g73970v3 (SULFUR DEFICIENCY-INDUCED 2) among others (Table 1, Fig-7). Genes in the dark red module were upregulated under moderate and high nitrate conditions but suppressed under low nitrate conditions and across ammonium treatments (Fig-7A-B). Positive module-trait correlations were observed for cortex, stele, and root cross-sectional areas and root hair density, while there was a strong negative correlation with total root length.

The yellow module contained the genes BRADI_2g12580v3 (cysteine-rich receptor-like protein kinase 15) and BRADI_3g55120v3 (PHOSPHATE STARVATION RESPONSE 2), which were positively correlated with N content in the whole plant and were highly expressed in the presence of higher N concentrations under both ammonium and nitrate applications (Fig-7). Both genes responded more strongly to nitrate application (downregulated in comparison with ammonium application), however, BRADI_2g12580v3 was significantly suppressed under low N conditions (Supplementary data 3). Conversely, the brown module contained BRADI_1g67440v3 (FAF-like, chloroplastic) and BRADI_4g02925v3 (protein JINGUBANG), which were negatively correlated with N content where their expression was suppressed under high N concentrations, and this effect was exacerbated in response to ammonium compared to nitrate (Fig-7).

## Discussion

### Strong associations between root phenotypes and transcriptomic responses suggest distinct transcriptional programs across traits

Root architecture and anatomical traits, which are controlled by many genes, exhibit significant phenotypic plasticity across different environments. They can adapt and change their structure in response to varying environmental conditions (Bray & Topp, 2018). These traits are influenced by genetic and environmental factors, allowing the plant’s roots to adapt and modify their structure depending on environmental conditions. Despite varying responses to N application among crop species, many of these traits have a conserved transcriptional response, suggesting a common genetic basis for adaptation to unfavorable conditions (Rogers & Benfey, 2015). We observed that root responses to ammonium and nitrate vary depending on their concentration and suggest that these responses are part of a nutrient foraging strategy to maximize root uptake area while minimizing the root system’s metabolic costs and preventing excessive nutrient accumulation.

Our study identified two key genes, BRADI_1g42930v3, which encodes an aspartyl protease family protein (APs), and BRADI_1g39167v3, which encodes phosphoenolpyruvate carboxylase 1 (PEPC 1), whose transcript levels correlated with the expression of numerous genes involved in root anatomical and morphological responses across varying concentrations of ammonium and nitrate (Fig-8). These two genes showed high betweenness centrality score among all hub genes (Supplementary data 5). A high degree betweenness centrality score indicates that a node has more connections with other nodes than the average (Kumar & Mukhtar, 2023). These genes potentially contribute to the optimization of metabolic costs by regulating root growth patterns that are responsive to nutrient availability. Supporting this, previous functional studies have highlighted the role of PEP carboxylase 1 in root architecture and nitrogen metabolism. Overexpression of PEPC 1 in rice demonstrated its influence on root system architecture, where a smaller root system formed including shorter primary roots at the seedling stage, as a response to altered metabolic fluxes (Lian et al., 2021). These changes were closely associated with adjustments in N assimilation and the expression of genes linked to the TCA cycle and glycolysis, highlighting the role of PEPC 1 in balancing nutrient uptake and metabolic energy costs. This evidence underscores the potential of PEPC 1 as a key player in optimizing root growth and nutrient use efficiency in response to environmental conditions.

**Figure 8:**
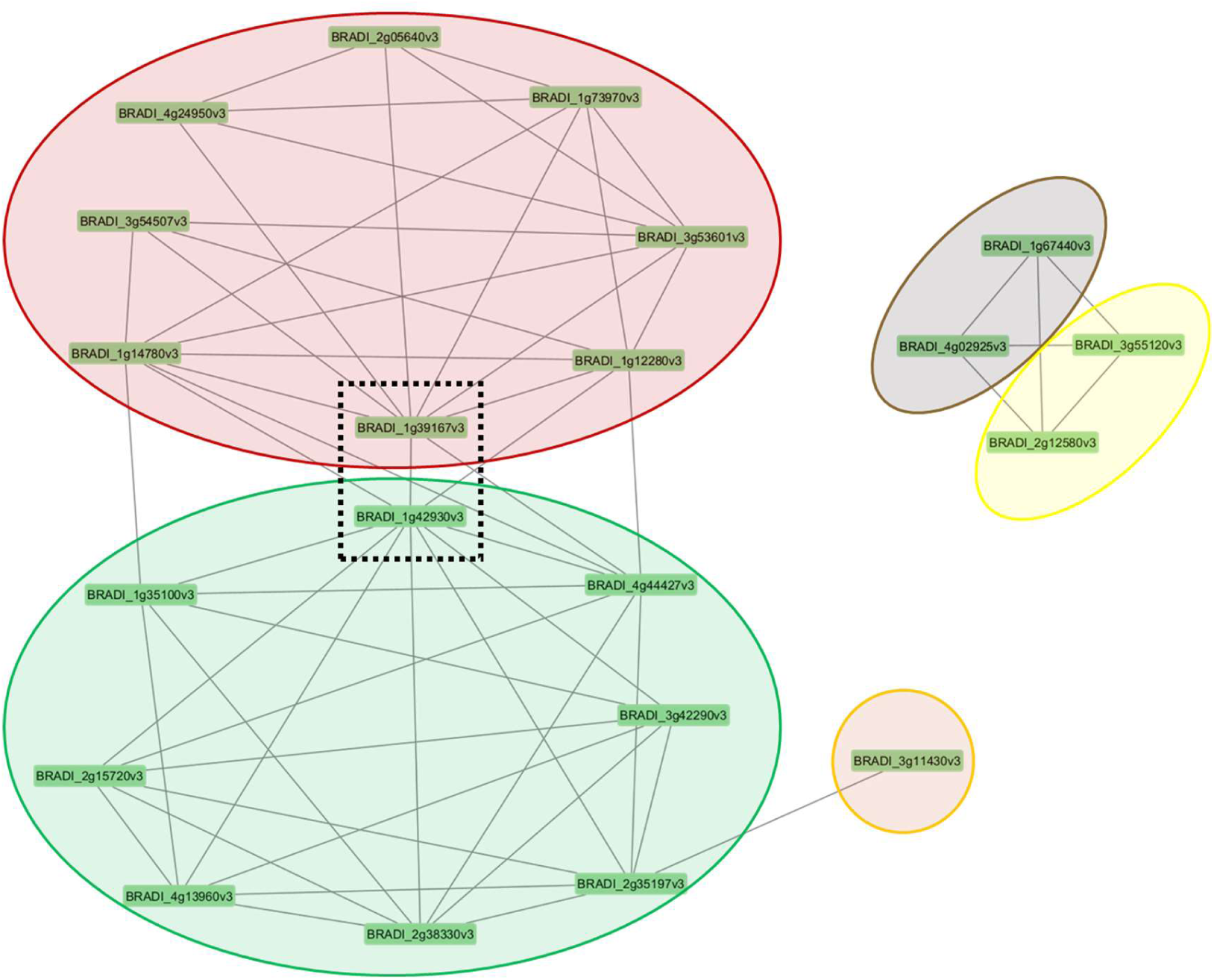
Clusters of co-regulated genes in *Brachypodium distachyon* under different nitrogen conditions. The network diagram displays modules of co-expressed genes, represented by colored ovals, with each node corresponding to an individual gene. The lines between genes indicate significant co-expression relationships calculated using topological overlap matrix. The central genes within the dashed box (BRADI_1g39167v3 and BRADI_1g42930v3) appear to link the two major modules (red and green), suggesting they may play a key regulatory role in connecting these gene networks.

On the other hand, an aspartyl protease family protein (ASPR 1) was shown to play a critical role in lateral root formation in *Arabidopsis thaliana* (Soares et al., 2019). Loss of function mutations in ASPR 1 led to a significant reduction in lateral root density, while it’s expression enhanced root branching. These findings suggest that expression of APs also have a role in controlling root system architecture to optimize nutrient uptake and minimize metabolic costs. The involvement of APs in processes like lateral root initiation further supports their role in nutrient foraging strategies, complementing the activities of PEPC 1 and its regulation of nitrogen uptake and metabolic efficiency.

### An aspartyl protease (APs) gene strongly correlated with root architecture response to low N availability

The brachypodium APs gene was strongly correlated with larger root system and lateral root development response under low N availability. Increasing the concentrations of nitrate and ammonium resulted in a smaller root system, characterized by reduced total root length and lateral root length. These traits are important for maximizing soil exploration and accessing N distributed in the soil profile (Lynch, 2019; Rogers & Benfey, 2015). Previous studies in Arabidopsis and maize have demonstrated similar results that the root foraging response to N deficiency involves the production of a large root system by elongation of both primary and lateral roots (Gao et al., 2014; Jia et al., 2019; Tian et al., 2005). A larger root system with a greater number of lateral branches increases the spatial extent of the root system, thereby enhancing the ability to acquire N from heterogeneous soils (Guo et al., 2014). However, some studies have shown an opposite response where N deficiency reduces root system area by decreasing lateral root density in cereals, including maize and rice (Ravazzolo et al., 2019). In rice, N deficiency modulated auxin levels, producing a smaller root system due to reduced number of lateral roots (Sun et al., 2019). The contrasting results of the two latter studies with the previous studies and our own results are likely due to different experimental conditions and sampling stages. For instance, rice studies used a half-strength growth medium containing only ammonium as N source, while we used an N-free medium as the lowest N concentration. Additionally, after two weeks, the N concentration in the brachypodium plants matched that of the non-germinated seeds, indicating that the plants primarily relied on their seed reserves during this early growth stage.

Auxin is a key regulator of lateral root development, influencing various stages from initiation to emergence. It promotes the formation of lateral root primordia by modulating cell division and expansion in the pericycle cells of the root (Du & Scheres, 2017). Our co-expression network analysis showed BRADI_1g35100v3 which encodes the auxin-responsive protein SAUR32 was co-expressed with the APs gene (BRADI_1g42930v3) (Fig-8). We reported that the expression of auxin-responsive protein SAUR32 (BRADI_1g35100v3) was negatively correlated with root cross-sectional area and root hair density (Fig-7, Fig-2A, Fig-3A). In Arabidopsis, it has been reported based on aspr1 mutants that this gene is involved in the deregulation of proteins associated with reactive oxygen species and auxin homeostasis, which is known to affect the number of lateral roots (Soares et al., 2019). For instance, APs protein family regulates auxin signaling by inhibiting the activity of AUXIN RESPONSIVE FACTORS (ARFs) during root growth (Cui et al., 2024; Rogg & Bartel, 2001; Soares et al., 2019; Su et al., 2020; Vierstra, 2009). In low N concentrations APs gene could potentially influence lateral root development indirectly by modulating AUXIN RESPONSIVE FACTORS (ARFs), leading to either a reduction or an increase in lateral root growth. Under normal nutrient conditions, expression of the BRADI_1g42930v3 homolog, *ASPR1*, in Arabidopsis led to reduced primary root growth and inhibited lateral root development (Soares et al., 2019), indicating its functional role in root formation. However, under N deficiency, no significant reduction in lateral root length was observed (Soares et al., 2019). Notably, in that study, the transcription level of *ASPR1* was not quantitatively assessed. In contrast, our results showed that ASPR1 suppression was correlated with the application of higher N concentrations, resulting in a reduced root system size and shorter lateral roots (Fig. 7, Fig. 1C). A recent genome-wide association study in maize highlighted the role of the GRMZM2G468657 gene, another homologue of APs in brachypodium (BRADI_1g42930v3), in root development and response to N deficiency (both its expression and polymorphisms) (Liu et al., 2023). To date, these two studies are the only ones that explored the role of the aspartic protease gene family in root architecture formation.

The brachypodium APs gene was also correlated with root structural changes at the anatomical level. Under ammonium and low nitrate application, APs gene expression was upregulated and associated with reduced root hair density and thinner roots. Root hairs significantly enhance the absorptive surface area, playing a crucial role in capturing immobile forms of N, such as ammonium (Zhiyong et al., 2022). The cross-sectional area, including the stele, influence the storage, transport, and metabolic efficiency of N absorbed from the soil (Burton et al., 2015). Larger stele areas can enhance vascular transport capacity, thereby improving the overall efficiency of nutrient transport to aboveground plant parts (J. Li et al., 2023; Lynch et al., 2021; Wang et al., 2019). It is reported that APs are also involved in programmed cell death in Arabidopsis and rice flowers (Niu et al., 2013; Phan et al., 2011). In roots, programmed cell death plays a crucial role in aerenchyma formation (Drew et al., 2000). It could highlight the potential roles of APs in root aerenchyma formation in higher N concentrations by modulating the programmed cell death in the root cortex area.

The APs gene (BRADI_1g42930v3) emerges as a potential regulator of root development and N response in brachypodium, interacting within complex biological networks involving hormone signaling and root architecture. Our results indicate its co-expression with ten other genes across different modules (Fig-8), suggesting functional relationships and coordination. Previous studies proposed that atypical aspartic proteases may have regulatory roles beyond housekeeping functions (Simoes & Faro, 2004). We showed that APs gene (BRADI_1g42930v3) could potentially modulate the root hormones as it was correlated to the expression of the BRADI_1g14780v3, which encodes a LOB domain-containing protein 37 correlated with root hair density and SAUR gene (BRADI_1g35100v3). Additionally, its co-expression with uncharacterized genes BRADI_1g12280v3 and BRADI_3g53601v3 further supports its role in N response and root formation. Interestingly, its transcript levels were associated with a PEPC 1 homolog (BRADI_1g39167v3) and exhibited contrasting correlations with various root phenotypes, highlighting its multifaceted regulatory potential.

### Reduction of metabolic costs by PEP carboxylase in response to higher N availability

The negative correlation of PEP carboxylase 1 (BRADI_1g39167v3), gene involved in carbon/nitrogen metabolism with mean length of lateral roots and total root length suggesting the adjustments at transcription scale to reduce metabolic costs associated with roots growth under N supply conditions. PEP carboxylase 1 plays an anaplerotic role in replenishing the TCA cycle intermediates and is involved in the plant’s response to stress conditions (Feria et al., 2016; Prinsi & Espen, 2018; Shi et al., 2015). The reduced expression of PEP carboxylase 1 and APs, along with the role of co-expressed genes such as SAUR and Aquaporin indicate a strategy to maintain efficient N uptake while reducing energy demands, thereby achieving a functional balance between root and shoot growth. Specifically, the downregulation of PEP carboxylase 1 and APs suggests that energy-intensive processes are being minimized. Meanwhile, SAUR genes, which are often involved in auxin signaling, could be playing a role in adjusting root growth patterns to optimize resource allocation. Aquaporins, which facilitate water and nutrient movement, help maintain proper nutrient transport with minimal energy expenditure. Thus, these adjustments allow the plant to conserve energy by modulating root activity, ensuring that N uptake is balanced with the energy needs of both root and shoot growth, maintaining a functional balance between these two systems.

When N is abundant in the environment, especially in the form of NO_3_ ^-^, we observe smaller root systems and increased shoot biomass. While the molecular signaling mechanisms that regulate root:shoot growth in response to N have been examined previously (David et al., 2019; Poitout et al., 2018), our study uncovered different morphological strategies by which brachypodium maintains the root:shoot ratio in response to different forms of N. Under high ammonium concentrations plants produce a smaller root system but maintain uptake area through increased lateral root formation (Fig-2), while under high nitrate concentrations plants invest in dense root hairs, which have lower energy costs, to maintain N uptake within the smaller root system (Fig-1). Both strategies maintain a functional equilibrium between belowground and aboveground resource acquisition (Z. Li et al., 2023; van Noordwijk et al., 1998).

Ammonium and nitrate have distinct assimilation pathways in plants with different energy requirements (Zayed et al., 2023). Specifically, nitrate assimilation is characterized by energy-intensive reduction steps involving the increased use of NAD(P)H, a byproduct of the TCA cycle, compared to NH_4_^-^ assimilation (Foyer et al., 2011; Liu & von Wirén, 2017). In our study, the expression of PEPC 1 remains constant across varying ammonium concentrations but increases with increasing nitrate concentration (Fig-7), which likely contributes to the higher energetic demands of NO_3_^-^ assimilation. In maize, increased levels of PEPC 1, along with phosphoglycerate mutase, suggested an escalation in respiratory metabolism in roots (Prinsi & Espen, 2018). Likewise, the Arabidopsis double mutant ppc1/ppc2 had severely reduced ammonium and nitrate assimilation under low N concentrations and accumulated more starch and sucrose than wild-type plants grown under nutrient sufficient conditions, suggesting overall decreased energy flux for N assimilation (Shi et al., 2015), which supports our results.

### Root growth suppression and aerenchyma formation avoids excessive N accumulation in high N environments

In addition to adjusting enzyme activity to reduce the metabolic cost of nitrogen uptake, plants modify root structures at various scales to balance N uptake with energy utilization. At the anatomical scale, under high ammonium concentrations brachypodium adapts their root energy utilization strategy by reducing the number of living cortical cells and forming aerenchyma in the root cortex. High concentration of ammonium is associated with ethylene production which induces aerenchyma formation (Zhu et al., 2010). However, the observed aerenchyma formation may also be attributed to the expression of the APs gene and the programmed cell wall death that results in downstream regulation of the cortex area. This explanation is in accordance with the observation of earlier research that atypical APs are involved in reproduction modulated programmed cell wall death in Arabidopsis and rice flowers (Ge et al., 2005; Huang et al., 2013).

On the other hand, the root aerenchyma formation could be an adaptive response not only to make an energy balance, rather a strategy to avoid excessive accumulation of N. Zhang et al. (2007) suggested that root growth suppression at high N concentrations is associated with elevated internal N levels, serving as a mechanism to prevent the excessive accumulation of N in shoot tissue. In our research, however, N concentrations in plant tissue remained relatively stable when media concentrations exceeded 1 mM ammonium or nitrate (Fig-1). We demonstrated that brachypodium avoids excessive accumulation not only by reducing the root system size but also by down regulating transporters, aquaporins, and possibly through senescence of cortical cells (Fig-5). Earlier studies observed decreased root aerenchyma formation in maize in response to either ammonium and nitrate fertilization (Konings & Verschuren, 1980). On the other hand, it is expected that the root aerenchyma formation could enhance plant growth in N limiting conditions (York et al., 2013). However, our study found that plants grown in high concentrations of ammonium produced aerenchyma in the middle part of their roots (Fig-3). This contrast findings in low N conditions could be due to early growth stage response of brachypodium roots to the application of high N concentration. This finding may suggest that aerenchyma formation is not only important for plants under low N conditions but may also play a role in nutrient acquisition under high ammonium conditions. Besides N uptake, this structural change accompanied with the strong transcriptional change could influence the uptake of other nutrients by the root system.

### N application influences gene expression and nutrient uptake beyond N excessive accumulation

Root anatomical modifications could potentially alter the uptake of nutrients that are otherwise adequately available. Our results highlighted the impact of N application on gene expression beyond those involved in N excessive accumulation. Notably, the transcript levels of BRADI_1g73970v3, which encodes the protein SULFUR DEFICIENCY-INDUCED 2, BRADI_2g38330v3, associated with zinc ion transmembrane transport, and BRADI_4g13960v3, a root-specific metal transporter, are affected by N conditions, with a correlation and potential direct influence on root anatomical changes. Our findings also underscore the complex relationship between sulfur and cysteine metabolism in response to ammonium and nitrate applications. Cysteine, being a key product of sulfur assimilation, plays a crucial role in incorporating sulfur into organic molecules, linking sulfur assimilation directly with N metabolism (Jobe et al., 2019). The upregulation of the sulfur deficiency gene (BRADI_1g73970v3) in higher N concentrations, particularly the moderated response under high ammonium concentrations, indicates a sophisticated sulfur deficiency response. Concurrently, cysteine synthesis, which is critical in plant immunity and cyanide detoxification, showed differential expression patterns depending on the N source, with significant implications for root hair development and plant defense mechanisms (Romero et al., 2014). These findings provide new insights into the complex interplay between nutrient uptake and root structure under varying N conditions, offering potential avenues for enhancing nutrient use efficiency in plants.

## Conclusion

Root growth adaptation to N supply is intricately dependent on both the dose and form of N, with distinct root architectural and anatomical responses observed for nitrate and ammonium. Our findings suggest that root growth and physiological responses to N form a concentration are driven by the plant’s need to maintain nutrient homeostasis while minimizing metabolic costs. Nitrate supply tends to suppress root growth, possibly as a strategy to optimize nutrient acquisition efficiency. In contrast, ammonium supply results in a more conservative approach, characterized by reduced cortex formation and decreased expression of transporters and aquaporins, reflecting a shift in resource allocation. This suggests that despite the diversity in crop types, a fundamental, evolutionarily conserved mechanism exists which governs how plants adapt their root structure and function to optimize nutrient acquisition in response to N availability. This dual response highlights the complex regulatory mechanisms plants employ to balance nutrient acquisition with the energetic constraints of high N availability, offering valuable insights for improving nutrient use efficiency in agricultural systems. Understanding these shared genetic pathways can provide valuable insights into breeding strategies aimed at enhancing N use efficiency across a wide range of crops, ultimately contributing to more sustainable agricultural practices.

## Material and Methods

### Growth conditions and treatments

*Brachypodium distachyon* is an ideal system for researching the impacts of nutrients in controlled environments due to its relatively small genome size and its adaptability to various substrates like soil or sand (Kellogg, 2015). We used brachypodium (Bd21-3) as model species to study root phenotypic plasticity and focused on seedlings to detect early N adaptive responses, rather than damage caused by prolonged stress. To compare root architectures, we grew the *Brachypodium distachyon* inbred Bd21-3, which is a relatively small model species that has a fully sequenced genome and an increasing number of genetic resources. Brachypodium seeds were sterilized for 5 minutes in 6% NaOCl + 0.1% Triton-X-100 under a clean bench and then washed five times with distilled, autoclaved water. The sterilized seeds were stored at 4°C on 1/3 Hogland medium containing nitrate as the N source for three days until germination at 18°C/16°C Day/night temperature in the dark. Two seedlings were then transferred to 250 ml plates with ammonium and nitrate applied as solo N form in 1/3 Hogland medium. The seedlings grew for up to 16 days using the grow screen agar system with 14 hours of light and 10 hours of dark. The experiment was conducted using a fully randomized block design with four replicates and 0, 0.18 mM, 0.37 mM, 0.75 mM, 1.5 mM, 3 mM, and 6 mM N concentrations. The experiment was repeated four times using an identical experimental setup. One repetition was used for the root phenotyping and root hair microscopy. The other repetitions were used for destructive measurements, such as N content, fresh weight (FW), and dry weight (DW) measurements. Missing values from each experiment were excluded, ensuring that at least two replicates were retained for analysis. The standard 1/3 Hogland medium includes KNO_3_, Ca (NO_3_) 2(H_2_O), MgSO_4_, KH_2_PO_4_, trace elements (MnCl_2_ · 4 H_2_O, CuSO_4_ · 5 H_2_O, ZnSO_4_ · 7 H_2_O, H_3_BO_3_, Na_2_MoO_4_ · 2 H_2_O), and [Fe ^+^-EDTA]. To apply ammonium and nitrate as solo N sources in the medium at different concentrations, we modified the standard solution by removing Ca (NO_3_) 2(H_2_O) for nitrate application and replacing KNO_3_ with (NH_4_) 2SO_4_ for ammonium application. To compensate for the absence of potassium and calcium from the modified medium, we added K_2_SO_4_ and CaCl_2_ to the 1/3 Hogland medium in the corresponding concentrations.

Our grow screen agar system was equipped with three high-throughput cameras that are able to take photos from the shoot front, above, and the root system separately (Suppl 10). The brachypodium seedlings were phenotyped every second day. The raw output of the phenotyping system was edited and cropped using Photoshop and then analyzed using the RhizoVision Explorer software (Seethepalli et al., 2021).

### Measurement of N concentration

To determine the N concentration in brachypodium seedlings, we harvested the 16-day old seedlings after germination and recorded their shoot and root fresh weights. After drying the shoot and root at 60°C for three days, we measured their dry weights. To analyze the N concentrations, we carefully ground the entire seedling, including the shoot and root, to a fine powder using a bead mill. We aliquoted 2 mg of this fine powder for analysis using a C/N macro elemental analyzer. By dividing the N concentration by the dry weight of the seedling, we were able to calculate the total N content with a high degree of accuracy.

### Root anatomical analysis

For our root anatomical analysis, we repeated the experiment under identical conditions and divided the root system into three equal sections corresponding to the top, middle, and bottom of the root system. We collected fresh root tissue from each divided section and preserved them in 75% ethanol for anatomical processing. To prepare the roots for sectioning, we solidified them in agarose and resized the blocks before gluing them onto a vibratome plate. We then used an Hm650v vibratome (Thermo Scientific Microm) to cut sections that were 40 μm thick. We collected three technical replicates for each section.

After collecting the individual sections with a fine brush, we transferred them to slides, humidified them with 1X phosphate-buffered saline, and observed them directly using confocal microscope (Nikon C2+) to acquire images of all the sections. We then analyzed the images using custom macros that we created with the open-source “PHIV Rootcell” toolset in ImageJ (Lartaud et al., 2015). The macros allowed us to trace the outlines of the cortex, stele, aerenchyma, vessels, and cells, and count the cell files. This careful quantification enabled us to determine the sizes of the cells and vessels.

### RNA extraction and transcriptome analysis

Brachypodium seeds were germinated on 1/3 modified Hogland medium with 1.5 mM nitrate for 3 days before being transferred to square petri dishes and grown for an additional 5 days. To obtain the transcriptome, nine brachypodium seedlings were grown in a modified 1/3 Hogland medium in square petri dishes in three replicates. Holes were made in the petri dishes to allow the shoots to grow outside. The roots of nine plants in each replicate were harvested, pooled together, and immediately frozen in liquid N.

The frozen root tissue was ground into a fine powder using a homogenizer in liquid N. Next, 450 µL of RTL buffer containing 10 µL of beta-mercaptoethanol was added to the tube, and the tissue was homogenized using a tissue homogenizer for 5 minutes. The samples were then transferred to an RNeasy Mini spin column placed in a 2 mL collection tube and centrifuged at 10,000 x g for 10 seconds. The filtrate was transferred to a new RNase-free tube, and 202 µL of absolute ethanol was added.

The extracted RNA was sequenced using Illumina technology, and sequencing libraries were prepared with the Ion Total RNA-Seq Kit v2 following the protocol from Life Technologies, USA. Quality assessment of the sequencing data was performed using FAST-QC. Cleaned reads were then mapped to the reference genome of *B. distachyon* (Brachypodium_distachyon_v3.0_genomic) using HISAT2 software (Schaarschmidt et al., 2020). Subsequently, the data were filtered using HTseq software to count how many aligned reads overlap with the exons of each gene. The edgeR package was utilized to identify differentially expressed genes, with a significance threshold of P value < 0.05 and a false discovery rate threshold of FDR < 0.1. Using a General Linear Model (GLM), we compared low versus moderate, low versus high, and moderate versus high concentrations of nitrate and ammonium. Gene ontology (GO) for biological processes were performed for DEGs using the “enricher” function of the “Biomart” library (Durinck et al., 2009) and visualized using clusterProfiler package in R (Yu et al., 2012).

### Weighted Gene Co-expression Network Analysis

Normalized RNA-seq data were analyzed using the WGCNA package in R to explore correlations between the transcriptome and the physiological phenotypes across the various treatments (Langfelder & Horvath, 2008). Genes with excessive missing values were excluded from the dataset and the remaining genes were used in the construction of weighted gene co-expression networks. Pearson correlation matrices for gene expression were computed for each network and transformed into connection strength matrices using an appropriate power function to ensure scale-free behavior. These connection strengths were then converted into a Topological Overlap Matrix (TOM).

TOM combined with average linkage hierarchical clustering was applied via the DynamicTree Cut algorithm to identify distinct co-expressed genes modules. To link these modules to phenotypic data, expression data was integrated with phenotypic responses across all N treatments. Modules with a correlation coefficient ≥ 0.4 and a p-value ≤0.05 were considered significant and selected for further analyses. Within each significant module, module membership (MM) and gene significance (GS) were calculated for all genes and the top genes were then identified based on their MM and GS for the phenotype of interest and were further investigated.

### Statistical analyses

Data were analyzed with R (R Core Team, 2013). Mean trait values were computed for each treatment, and the variation among the mean trait values of the two N treatments and concentrations was evaluated using analysis of variance (ANOVA). When the ANOVA indicated significance at the 5% probability level, the Tukey’s HSDtest was used to assess differences between treatments.

## Acknowledgements

We sincerely thank Dr. Robert Koller, Dr. Tobias Wojciechowski, and Prof. Uwe Rascher for their invaluable guidance as members of the project advisory board. We are also grateful to Andrea Neuwohner, Sabine Preiskowski, and Prof. Ingar Janzik from IBG-2 for providing lab space and support for RNA extraction from root systems. Special thanks to Kiran Suresh at the Institute for Cellular and Molecular Botany (IZMB), Bonn University, for assisting with root cross-sections and microscopic imaging. We are also thankful to Dr. Rizwan Riaz and the HOCBio Center at University of Illinois for providing cluster computing access and assistant in bioinformatics analysis.

## Funding

This work has been funded by the German Research Foundation under Germany’s Excellence Strategy, EXC-2070 - 390732324 – PhenoRob. Research reported in the publication was supported by the Foundation for Food & Agriculture Research under award number – Grant ID: 602757 (to AMC). The content of this publication is solely the responsibility of the authors and does not necessarily represent the official views of the Foundation for Food and Agriculture Research

## Author Contributions

HR conceived the study, designed and conducted lab experiments, performed computational analysis, and led manuscript preparation. DS contributed to the WGCNA and RNA-seq data analysis. CA contributed to the RNA-seq data analysis. BM contributed to manual annotation for root hair image analysis. LS supervised root anatomical analysis and microscopy. BS conceived the study, designed the experiments, supervised computational and bioinformatics data analysis, and contributed to manuscript preparation. AM-C conceived the study, designed the experiments, supervised computational and bioinformatics data analysis, and led manuscript preparation. All authors contributed to drafting the manuscript.

## Notes

### Competing Interest Statement

The authors have declared no competing interest.

### Summary of Updates

Figure 2b revised; author affiliations updated.

