## supplementary figures for "Regulatory Analysis of Root Architectural and Anatomical Adaptation to Nitrate and Ammonium in *Brachypodium distachyon*"

**Supplementary figure 1**

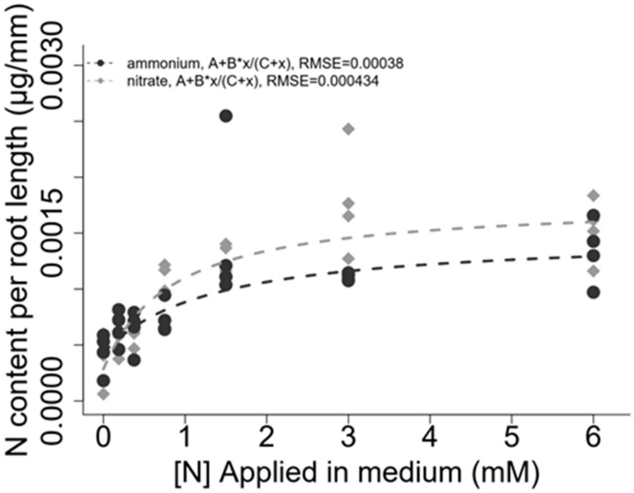


**Supplementary figure 2 and 3**

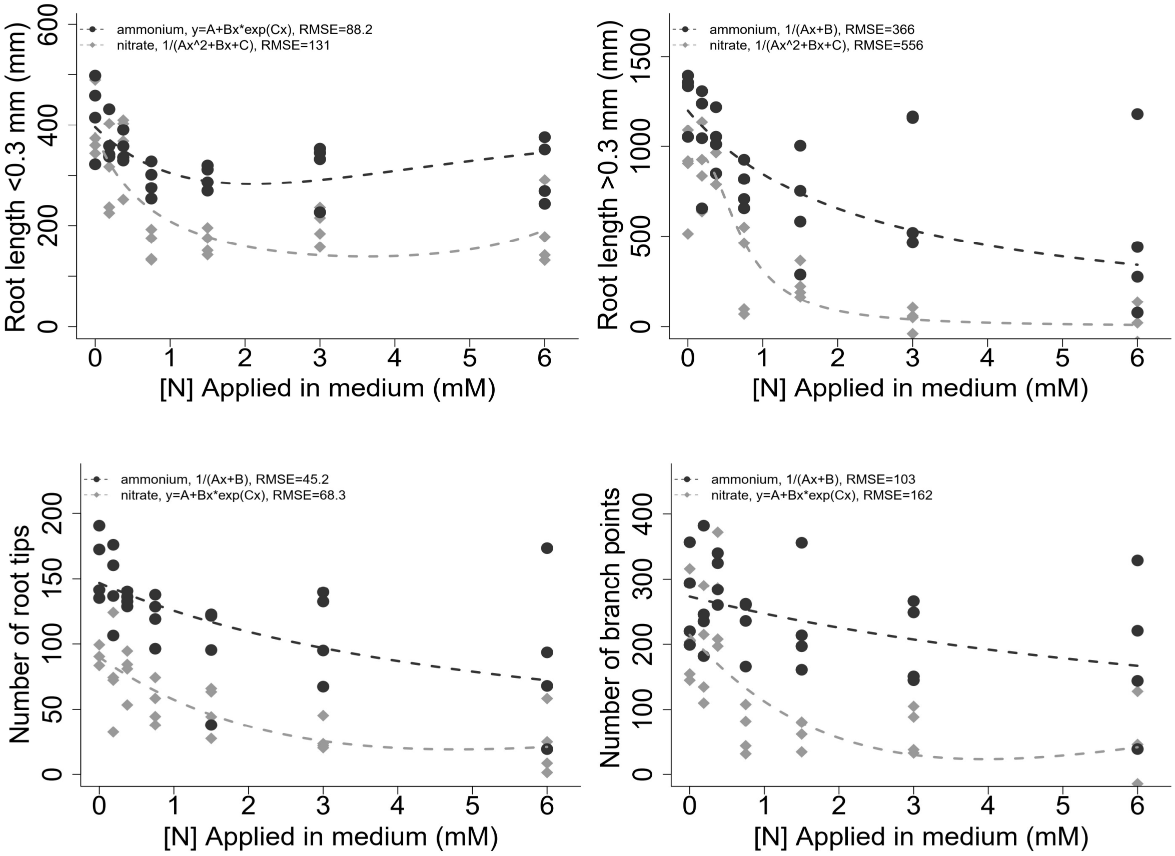

**Supplementary figure 4**

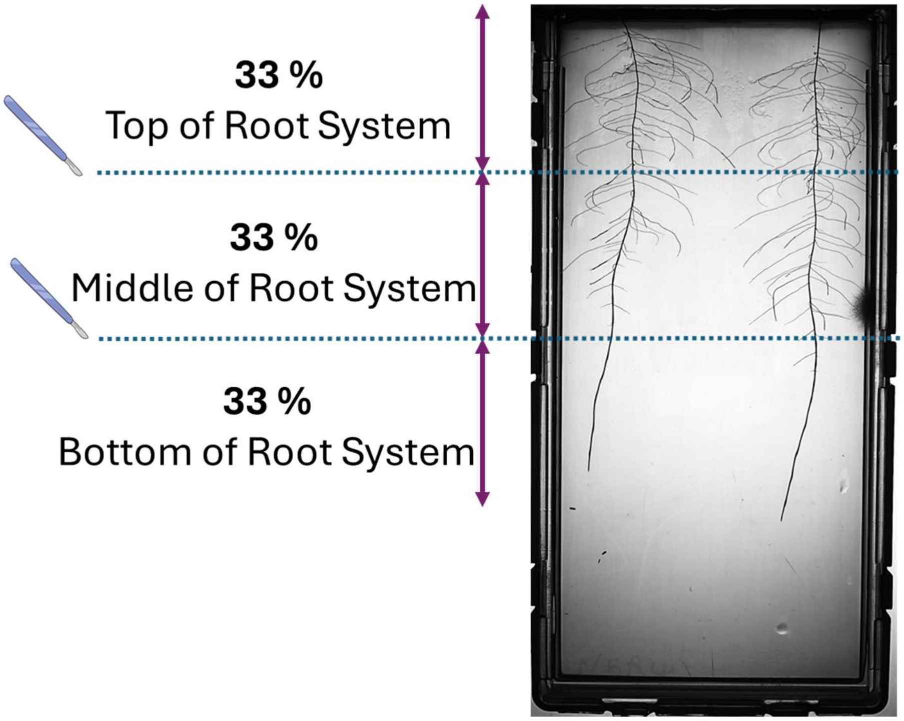


**Supplementary figure 5**

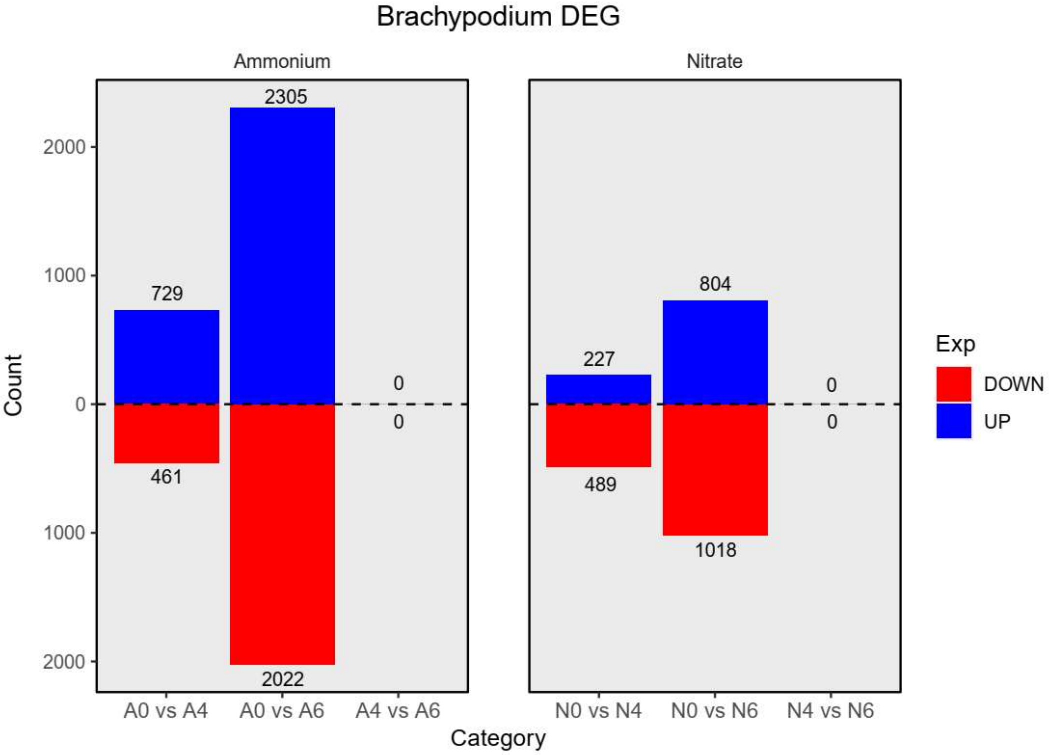

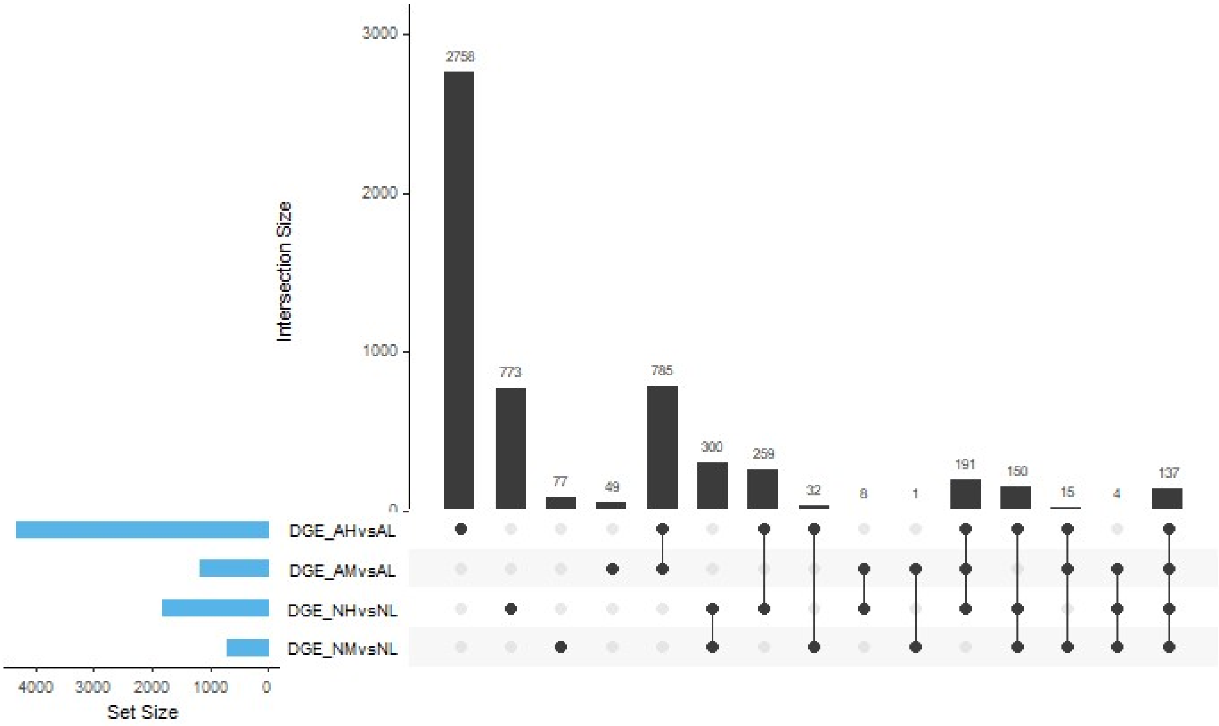


**Supplementary figure 6**

**GO enrichment of up regulated genes in moderate vs low nitrate**

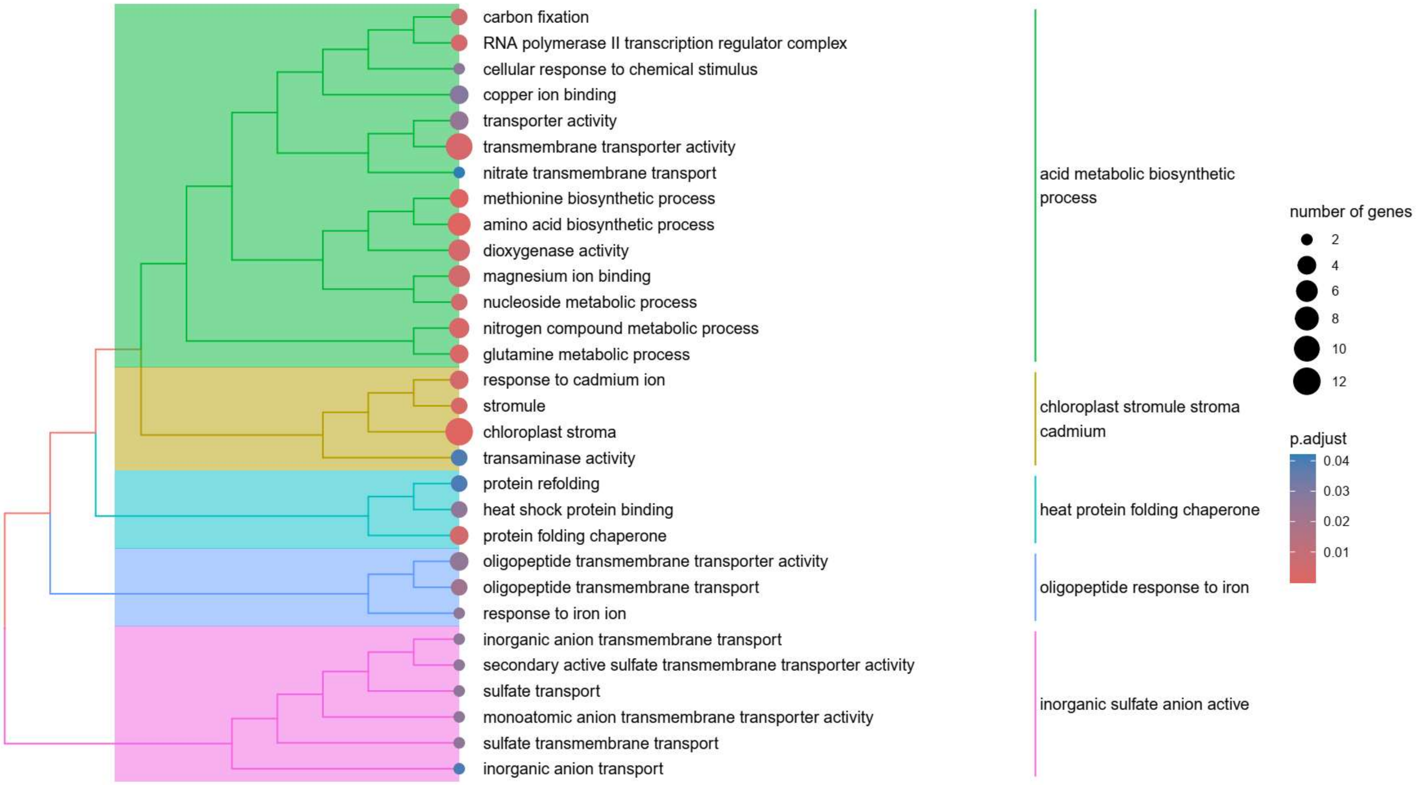

### GO enrichment of up regulated genes in high vs low nitrate

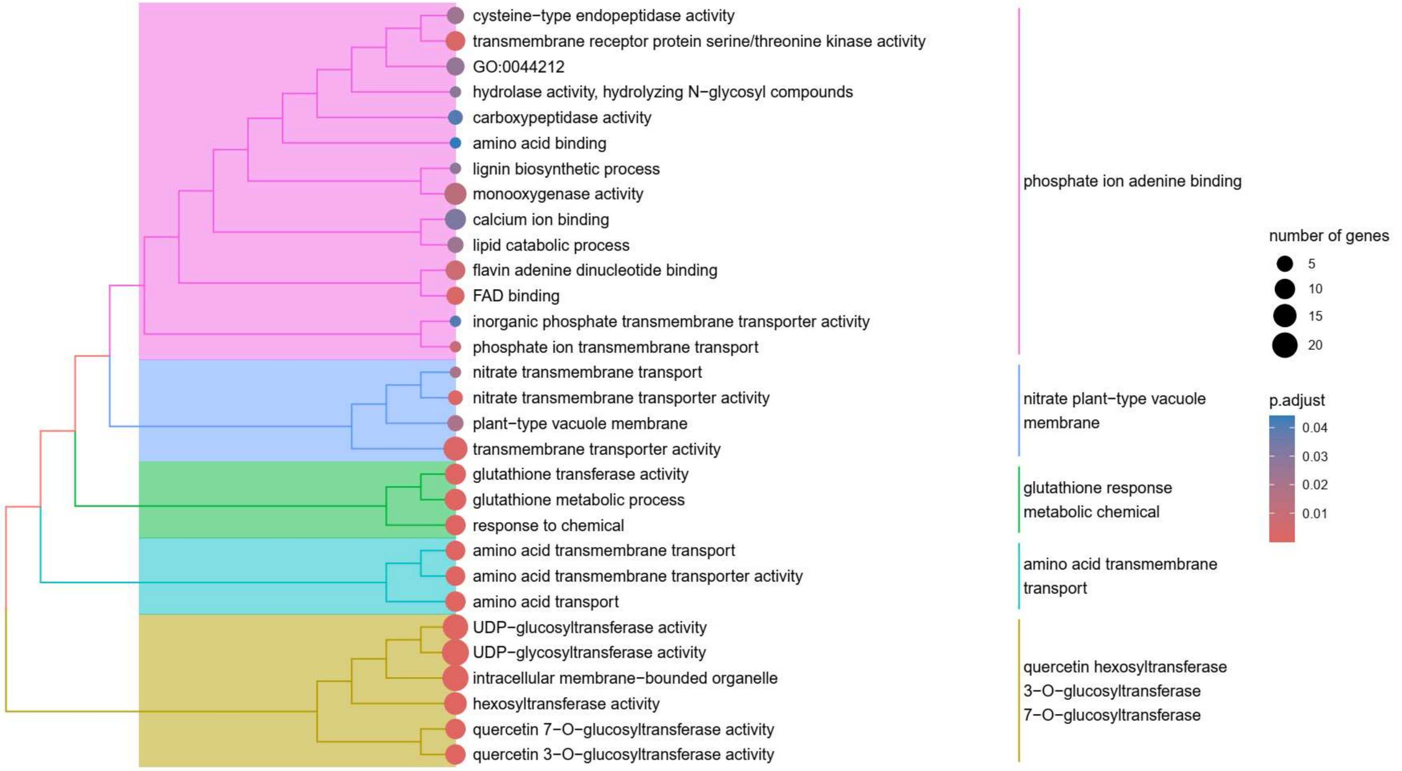


### GO enrichment of down regulated genes in moderate vs low nitrate

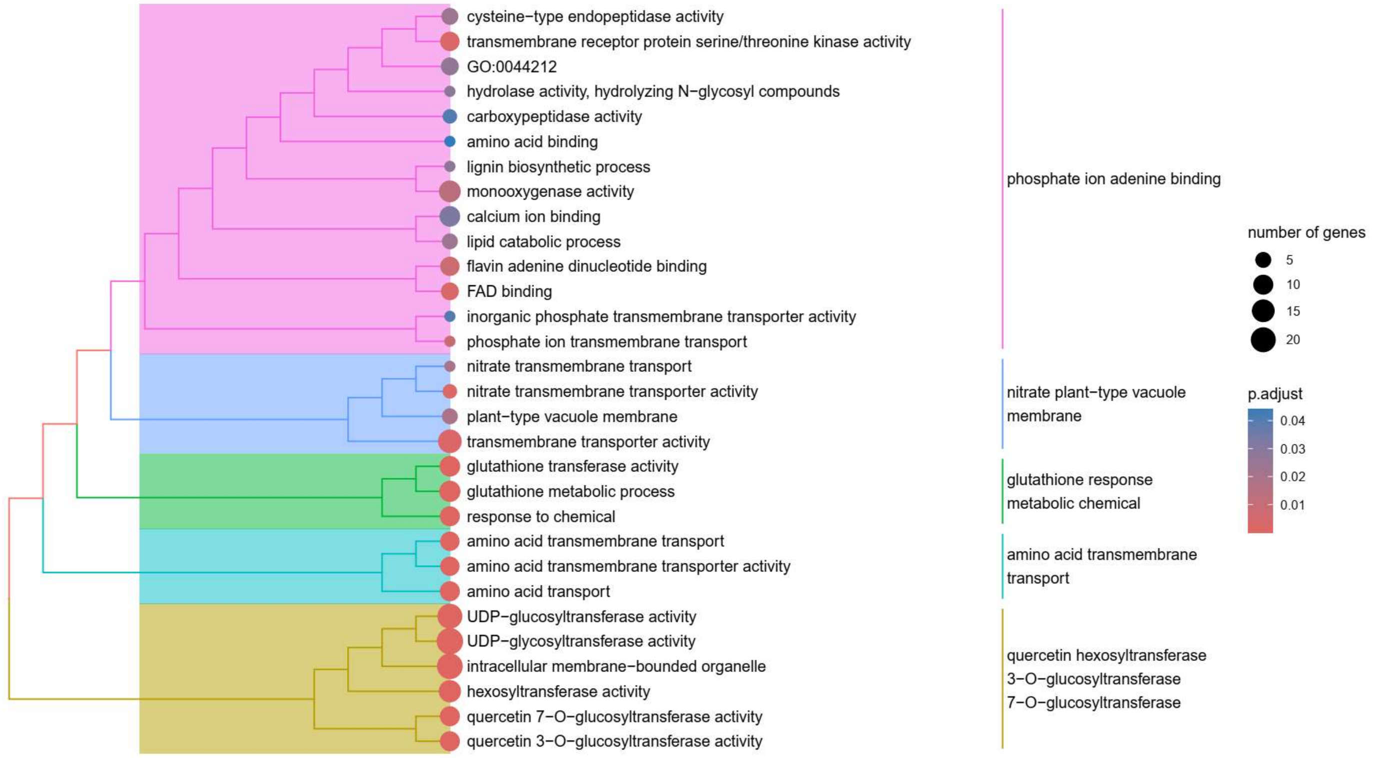

### GO enrichment of down regulated genes in high vs low nitrate

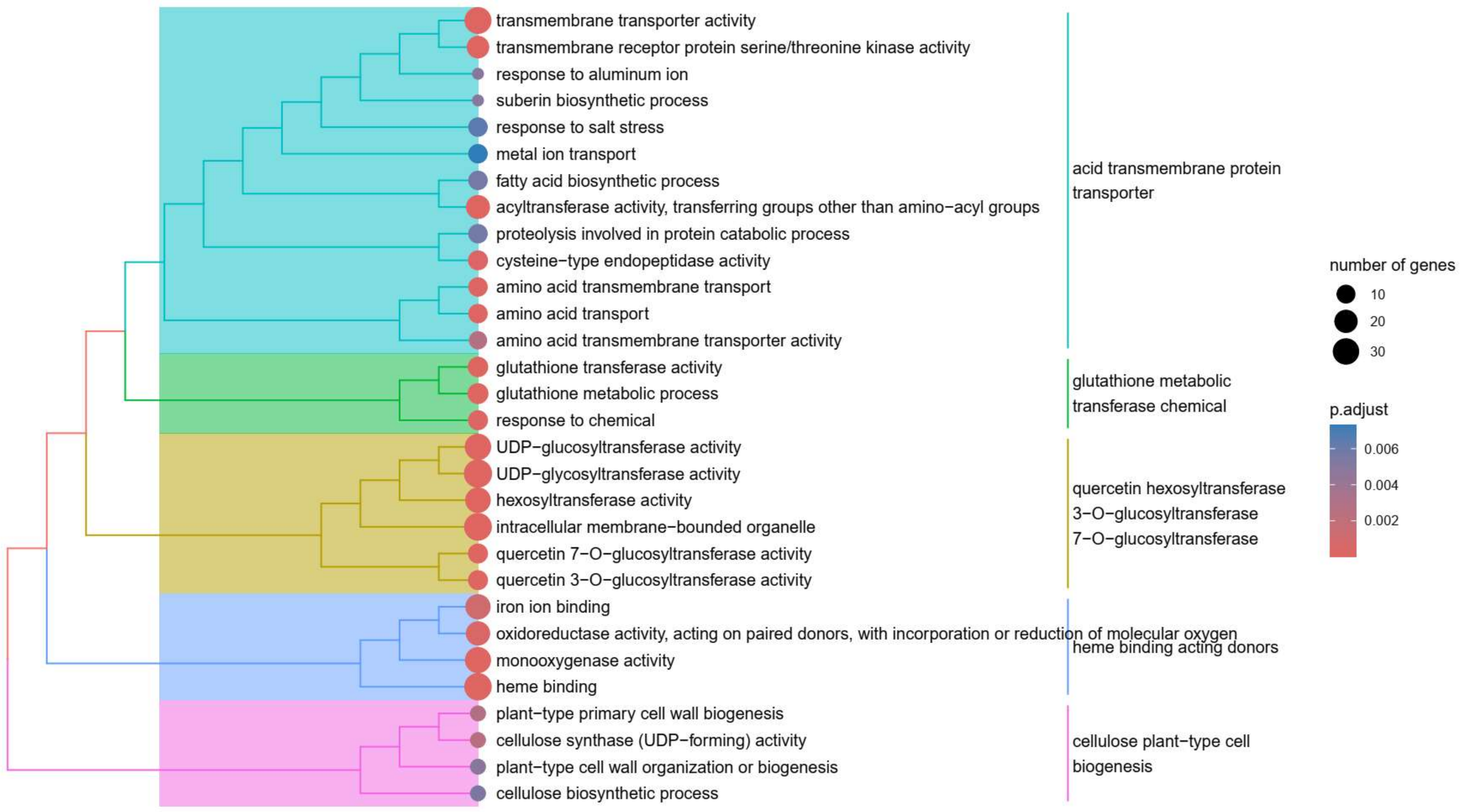


### Supplementary figure 7

### GO enrichment of up regulated genes in moderate vs low ammonium

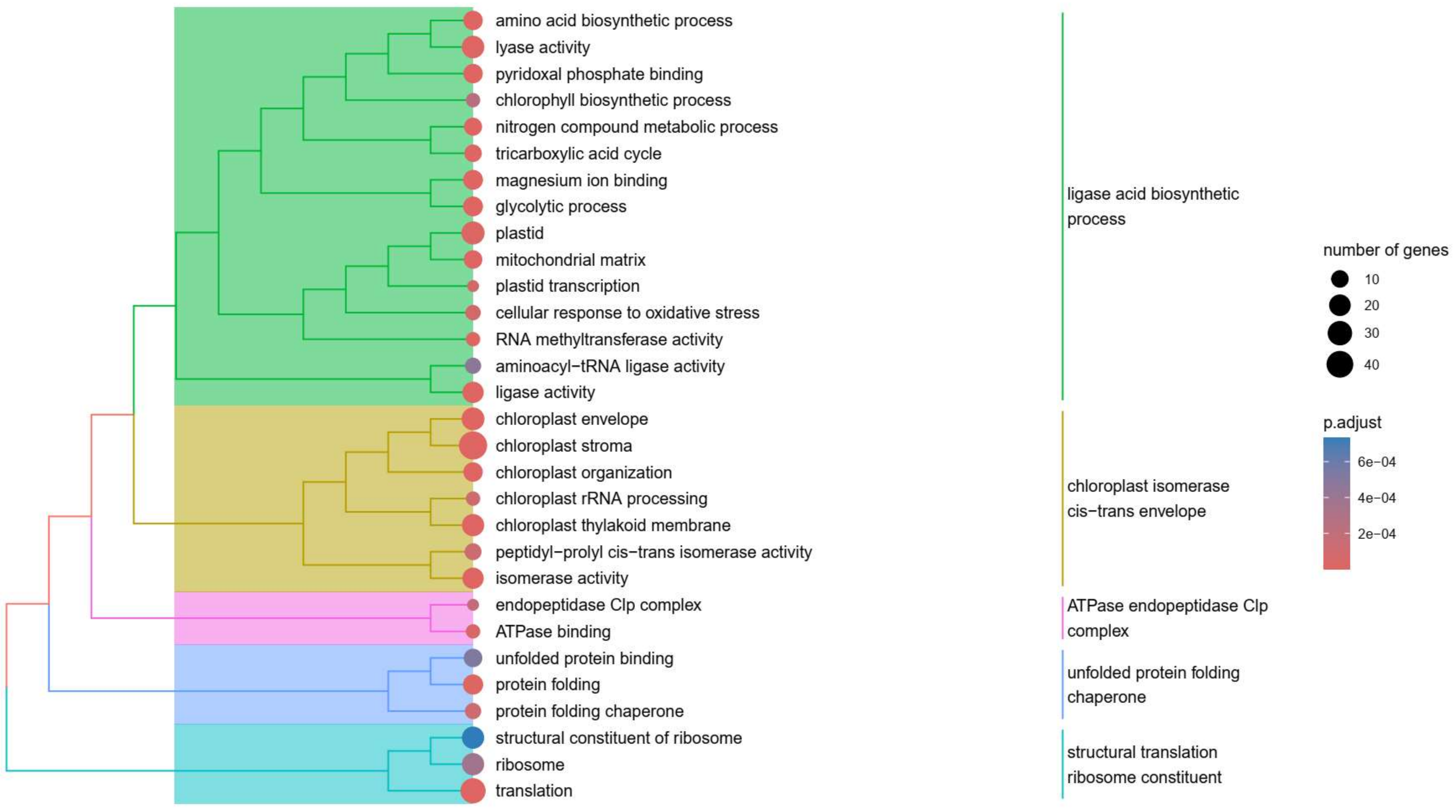

### GO enrichment of up regulated genes in high vs low ammonium

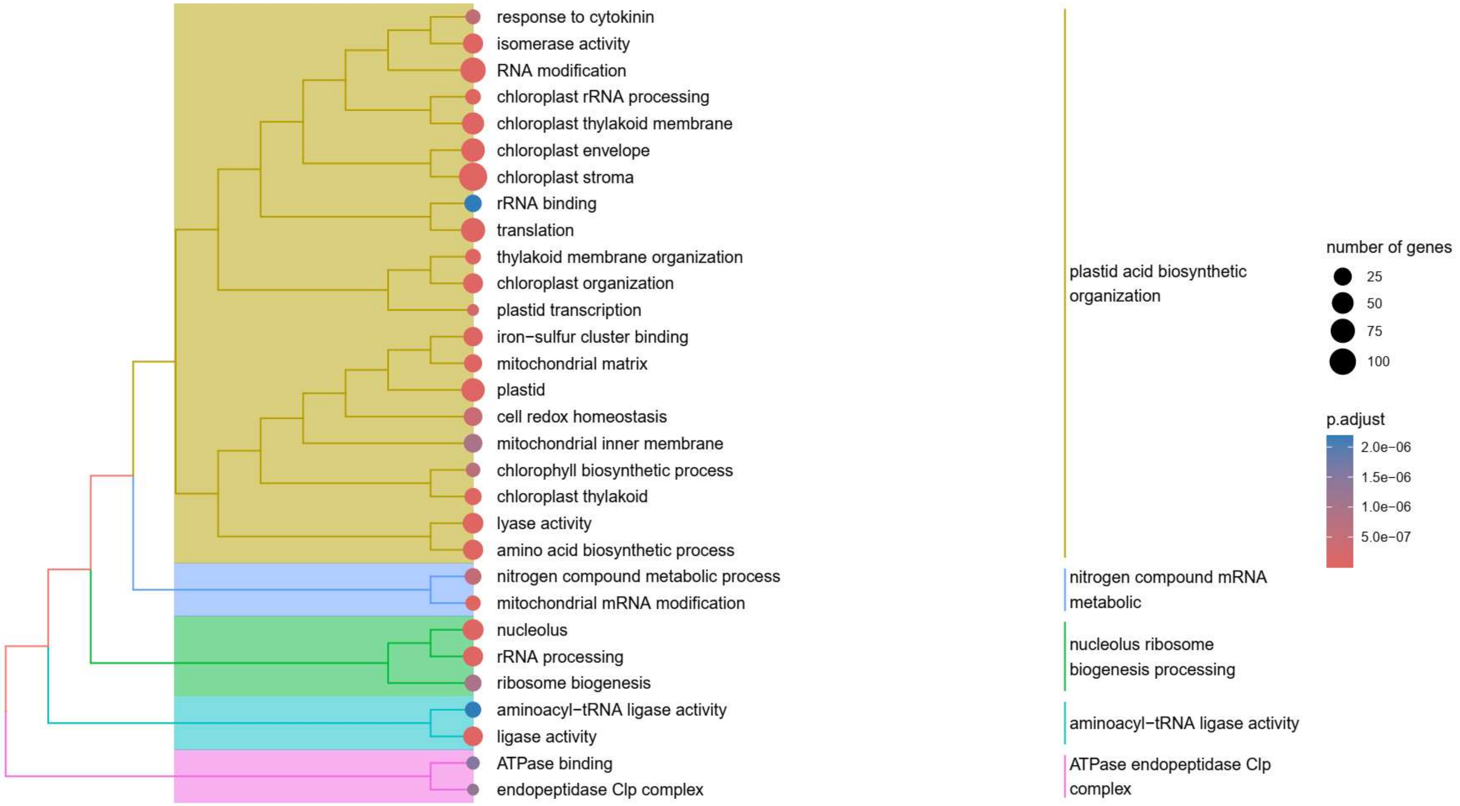


### GO enrichment of down regulated genes in moderate vs low ammonium

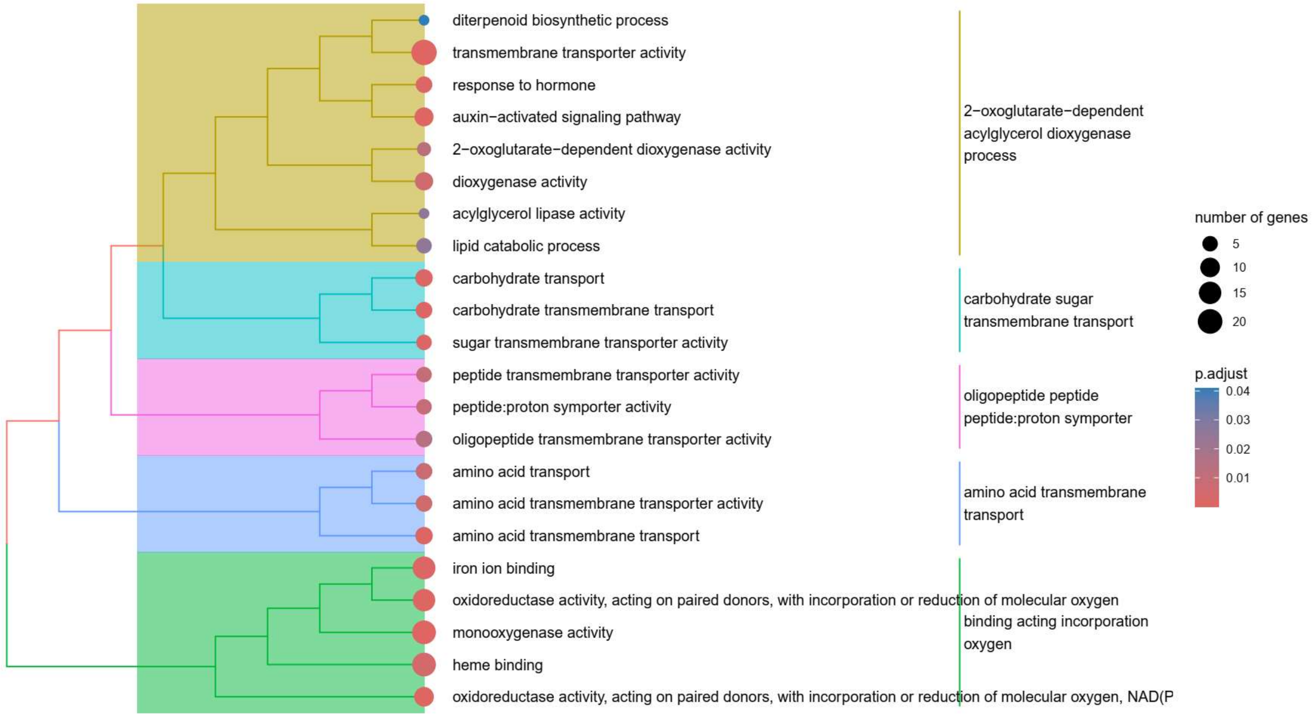

### GO enrichment of down regulated genes in high vs low ammonium

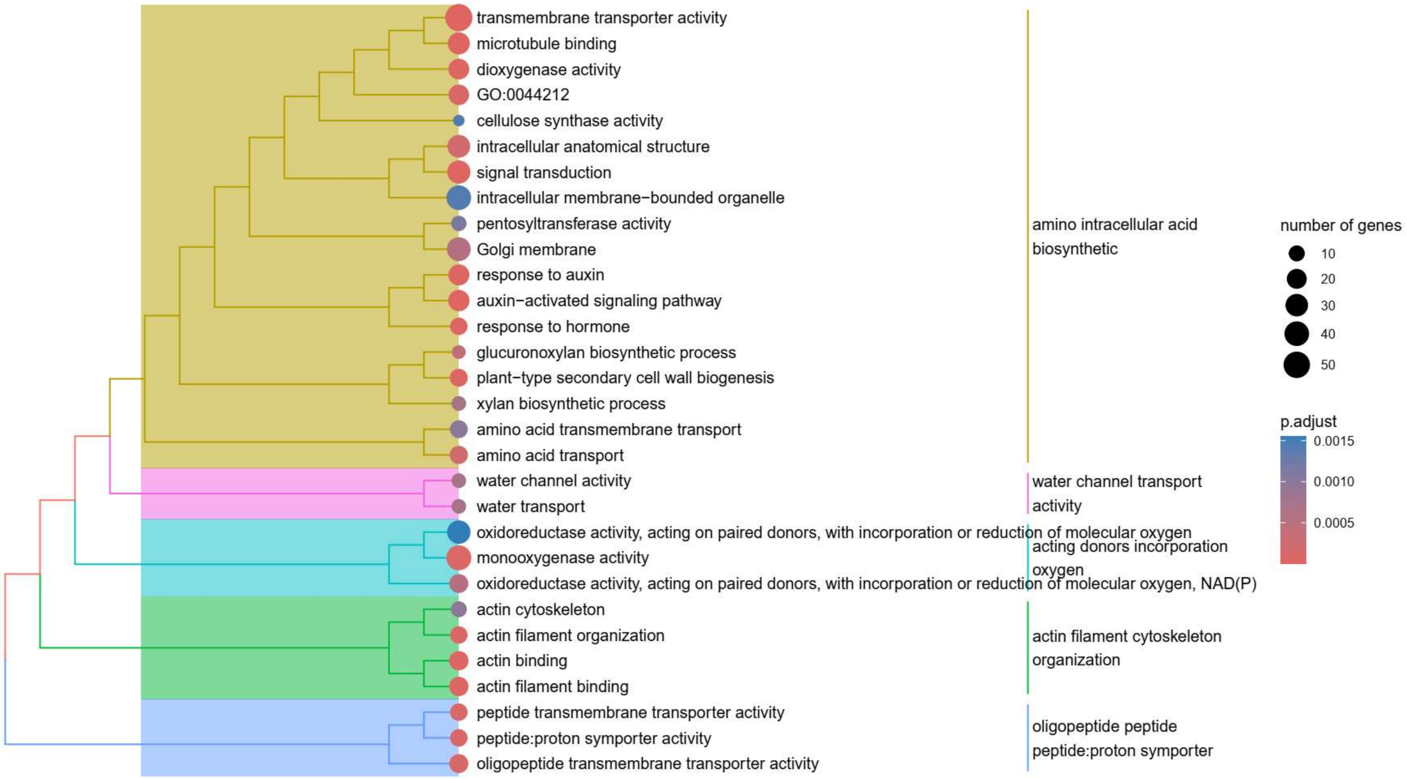


### Supplementary Table 8

| Name | Gene ID |
|---|---|
| NRT1.1 | BRADI_3g33040v3 |
| NRT1.2 | BRADI_1g37330v3 |
| NRT1.3 | BRADI_3g47010v3 |
| NRT1.4 | BRADI_2g41060v3 |
| NRT1.5 | BRADI_3g53380v3 |
| NRT2.1 | BRADI_3g01270v3 |
| NRT2.2 | BRADI_3g01250v3 |
| NRT2.3 | BRADI_3g01277v3 |
| NRT2.4 | BRADI_3g01290v3 |
| NRT2.5 | BRADI_2g47640v3 |
| NRT2.6 | BRADI_2g26210v3 |
| NRT2.7 | BRADI_2g40740v3 |
| NRT3.1 | BRADI_3g47710v3 |
| NRT3.2 | BRADI_3g47720v3 |

**Supplementary figure 9**

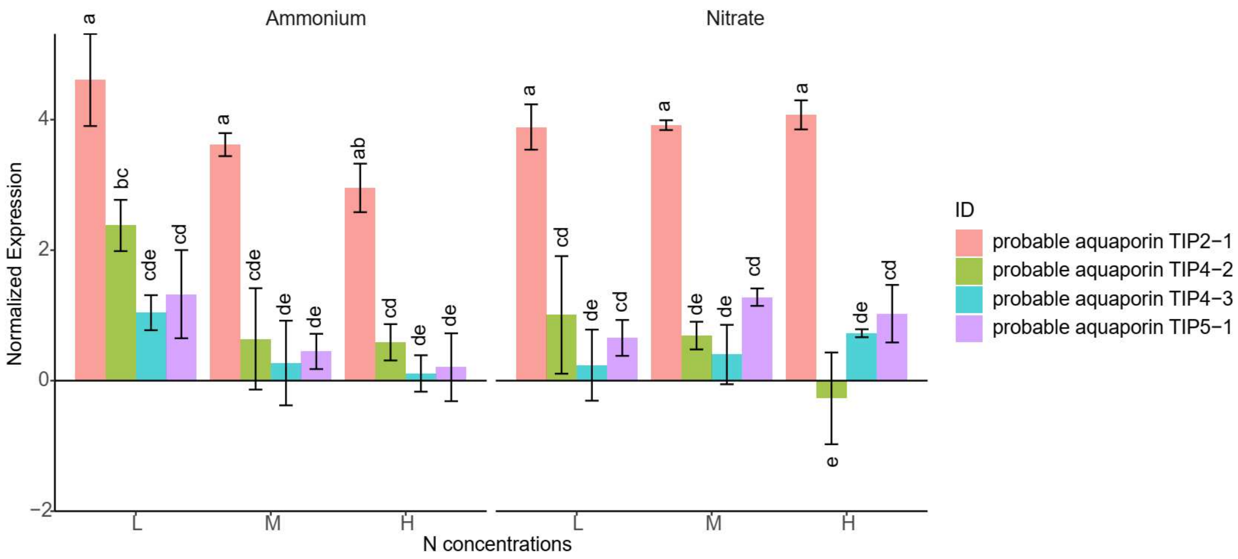

### Supplementary Table 10

| Gene ID | Name | Modules | Phenotype + correlation | Phenotype - correlation |
|---|---|---|---|---|
| BRADI_1g67440v3 | Protein FAF-like, chloroplastic [ Brachypodium distachyon (stiff brome) | Brown | | Nitrogen content |
| BRADI_4g02925v3 | Protein JINGUBANG [ Brachypodium distachyon (stiff brome) | Brown | | Nitrogen content |
| BRADI_1g14780v3 | LOB domain-containing protein 37 [ Brachypodium distachyon (stiff brome) ] | Dark red | Root hair density | Total root length |
| BRADI_3g53601v3 | Uncharacterized LOC104581291 [ Brachypodium distachyon (stiff brome) | Dark red | Root hair density | Total root length and mean of lateral root |
| BRADI_1g73970v3 | Protein SULFUR DEFICIENCY-INDUCED 2 [ Brachypodium distachyon (stiff brome) ] | Dark red | Cross-sectional area | |
| BRADI_1g39167v3 | Phosphoenolpyruvate carboxylase 1 [ Brachypodium distachyon (stiff brome) ] | Dark red | Cortex, stele, and root cross-sectional area, root hair density | Total root length |
| BRADI_1g12280v3 | Uncharacterized LOC104582693 [ Brachypodium distachyon (stiff brome | Dark red | Root hair density, cortex, stele, and root cross-sectional area | Total root length |
| BRADI_2g05640v3 | Clavaminate synthase-like protein At3g21360 [ Brachypodium distachyon (stiff brome) | Dark red | Cross-sectional and cortex area | |
| BRADI_4g24950v3 | Proline-rich protein haeiii subfamily 1 [ Brachypodium distachyon (stiff brome) | Dark red | Root cross-sectional and cortex area | |
| BRADI_3g54507v3 | Uncharacterized | Dark red | Root cross-sectional and cortex area | |
| BRADI_2g38330v3 | Involved_in zinc ion transmembrane transport | Lightgreen | Mean length of lateral roots and total root length | |
| BRADI_4g13960v3 | Root-specific metal transporter | Lightgreen | Total root length | |
| BRADI_1g42930v3 | Aspartyl protease family protein At5g10770 [ Brachypodium distachyon (stiff brome) ] | Lightgreen | Mean length of lateral roots and total root length | Root hair density, cortex and root cross-sectional area, stele area |
| BRADI_4g44427v3 | Probable serine/threonine-protein kinase WNK8 | Lightgreen | | Root cross-sectional and cortex area |
| BRADI_3g42290v3 | Protein DETOXIFICATION 33 | Lightgreen | | Cross-sectional and cortex area |
| BRADI_2g15720v3 | 3-ketoacyl-coa synthase 11 [ Brachypodium distachyon (stiff brome) | Lightgreen | | Root cross-sectional and cortex area |
| BRADI_2g35197v3 | Serine carboxypeptidase 2 [ Brachypodium distachyon (stiff brome) | Lightgreen | | Root cross-sectional and cortex area |
| BRADI_1g35100v3 | Auxin-responsive protein SAUR32 [ Brachypodium distachyon (stiff brome) ] | Lightgreen | | Cortex and root cross-sectional area, root hair density |
| BRADI_1g26990v3 | Expansin-like B1 [ Brachypodium distachyon (stiff brome) | Orange | Central metaxylem | |
| BRADI_3g11430v3 | Nicotianamine aminotransferase A [ Brachypodium distachyon (stiff brome) | Orange | Central metaxylem | |
| BRADI_2g12580v3 | Cysteine-rich receptor-like protein kinase 15 | Yellow | Nitrogen content | |
| BRADI_3g55120v3 | Protein PHOSPHATE STARVATION RESPONSE 2 | Yellow | Nitrogen content | |

**Supplementary figure 11**

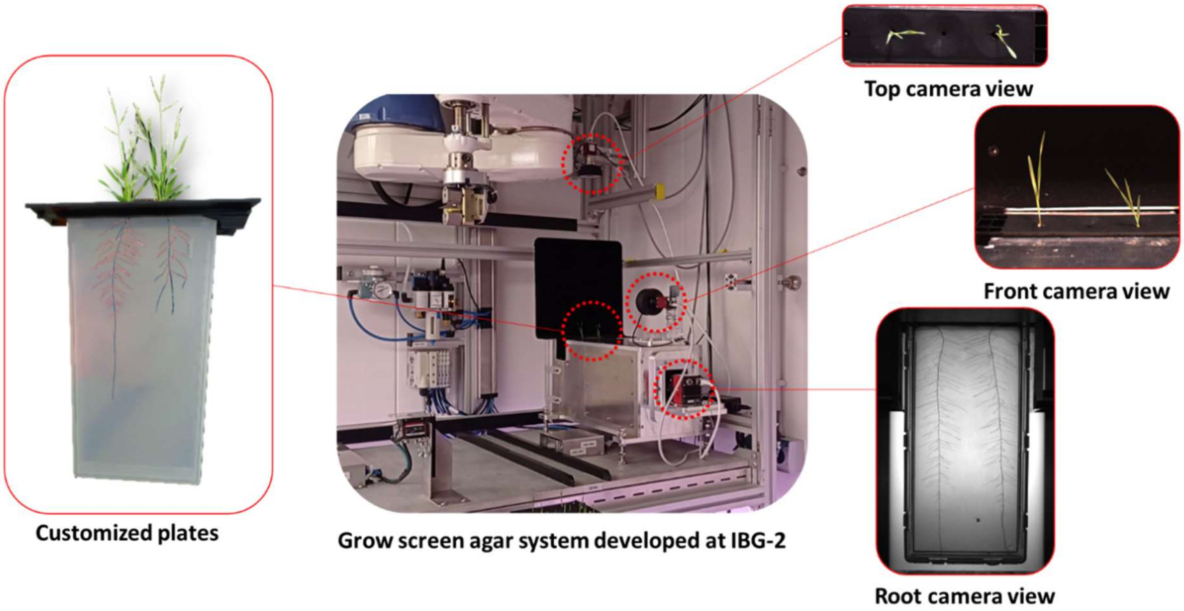
